# A divergent mutational and clonal landscape in aged HSCs is not linked to aging-associated clonal hematopoiesis

**DOI:** 10.64898/2026.01.24.701060

**Authors:** Maria Carolina Florian, Amanda Amoah, Kalpana J. Nattamai, Karin Soller, Angelika Vollmer, Markus Hoenicka, Andreas Liebold, Katharina Bauer, Jan-Phlipp Mallm, Hartmut Geiger, Medhanie A. Mulaw

**Affiliations:** Institute of Molecular Medicine and Stem Cell Aging, University of Ulm, Albert-Einstein-Allee 11, D-89081 Ulm, Germany; Division of Experimental Hematology and Cancer Biology, Cincinnati Children’s Hospital Medical Center and University of Cincinnati, Cincinnati, OH, USA; Department of Cardiothoracic and Vascular Surgery, Ulm University Hospital, Ulm, Germany; Program for Regenerative Medicine, IDIBELL, 08908 L’Hospitalet de Llobregat, Barcelona, Spain; DKFZ and BioQuant Center scOpenLab, BioQuant, Heidelberg, Germany; Division of Chromatin Networks, German Cancer Research Center (DKFZ), Heidelberg, Germany; Unit for Single-cell Genomics, Medical Faculty, University Hospital Ulm, Albert-Einstein-Allee 11, D-89081 Ulm, Germany

**Author notes:** Authors with equal contribution (Hartmut Geiger, Medhanie A. Mulaw). Corresponding author/Lead Contact: Medhanie A. Mulaw, Ph.D.

## Abstract

Somatic mutations accumulate throughout life and can serve as endogenous markers to trace cellular lineage relationships. In hematopoietic stem cells (HSCs), aging is associated with functional decline and clonal skewing, yet how somatic mutational histories shape clonal architecture at single-cell resolution remains incompletely understood. Here, we leverage single-cell RNA sequencing from murine and human long-term HSCs to identify expressed somatic single-nucleotide variants and reconstruct mutation-based genealogies. To address technical noise inherent to single-cell transcriptomes, we implement a stringent joint variant filtering strategy that exploits daughter cell relationships and probabilistic modeling to distinguish true somatic events from artifacts. Using these high-confidence variants, we infer phylogenetic and clonal relationships among individual HSCs and uncover striking age-dependent differences in lineage structure. Whereas young HSCs show little evidence of hierarchical clonal organization, aged HSCs frequently exhibit structured genealogies consistent with clonal expansion. Integration of mutational, transcriptional, and signature-based analyses further reveals that genetically distinct clones in aged samples are transcriptionally divergent and enriched for pathways linked to aging and DNA damage responses. Together, our findings demonstrate that somatic mutations recovered from single-cell transcriptomes can resolve HSC clonal evolution and reveal age-associated alterations in stem cell dynamics.

## Introduction

Stem cell exhaustion and genomic instability are regarded as hallmarks of aging (Lopez-Otin et al., 2013). In addition to cells acquiring DNA mutations with varying degrees of adaptive potential, these events lead to changes in heterogeneity and clonality among individual cells, which consequently contribute to aging-related diseases.

Hematopoietic cells generally show a modest increase in DNA mutations and elevated levels of DNA damage and replication stress (Beerman et al., 2014; Flach et al., 2014; Moehrle et al., 2015; Rossi et al., 2007; von Zglinicki et al., 2001). Aging correlates with an increased incidence of clonal hematopoiesis of indeterminate potential (CHIP), also termed age-related clonal hematopoiesis (aCH) (Jaiswal and Ebert, 2019; Steensma et al., 2015). aCH is considered a pre-disease stage that may evolve into myelodysplastic syndrome (MDS) or acute myeloid leukemia (AML). Specifically, aCH is defined as the gradual, clonal expansion of hematopoietic stem and progenitor cells (HSPCs) carrying specific, disruptive, and recurrent genetic variants in individuals without a clear diagnosis of hematological malignancies. Whether clonality begins within the stem cell pool during aging and whether this is linked to heterogeneity among HSCs remain matters of debate (Haas et al., 2018; Moehrle and Geiger, 2016; Moehrle et al., 2015; Nijnik et al., 2007), as the mutational load and types of mutations in individual HSCs are difficult to assess accurately, primarily due to experimental limitations in current single-cell sequencing approaches for determining mutational load in individual cells.

Single-cell sequencing still poses a technical challenge due to low coverage and high noise, which lead to a high false-positive rate of single-nucleotide variants (SNVs) or mutations (Lun et al., 2016). Novel computational filtering approaches have been designed to reduce the influence of confounding factors (Luquette et al., 2019). In this study, the authors used allele-specific amplification imbalance as a framework for statistical estimation of mutations in single cells by leveraging the allele frequencies of neighboring heterozygous single-nucleotide polymorphisms (SNPs). Additionally, single-cell multiple displacement amplification (SCMDA) coupled with single-variant cell variant caller (SC-caller) has been reported to significantly reduce false-positive call rates (Dong et al., 2017). More recently, mathematical modeling coupled with targeted DNA sequencing of frequently mutated genes in single cells was used to generate clonal hierarchies within leukemia samples (Miles et al., 2020). Despite these and other advances, current methods are not able to fully eliminate uncertainty in the process of correcting SNV calling, as they are still based on multiple predictions and juxtaposed genomic contexts. Additionally, current methods rely on DNA sequencing, and it is technically not possible to evaluate whether these mutations are expressed in a given cell, limiting the interpretation of the functional role of these mutations. Overall, SNV determination in single-cell data and linkage to gene expression thus remain major challenges in single-cell sequencing approaches (Lahnemann et al., 2020).

Due to current limitations in calling SNVs in single cells, including HSCs, comparisons of the actual SNV rate in young vs. aged HSCs and its relationship to clonal hematopoiesis during aging are lacking. We provide a simple and accurate approach to quantify SNVs in single-cell RNA-seq data from young and aged murine and human HSCs, enabling integrated interpretation of expression and SNV data from individual HSCs. Using this approach, we show that the overall aging-related accumulation of mutations in HSCs is modest and primarily linked to the chemical instability of DNA. In humans, mutations in genes known to be relevant to clonal hematopoiesis were identified in both young and aged HSCs at similar frequencies. Targeted sequencing of human bone marrow (BM) cells from the same individuals from which the HSCs were analyzed revealed that the mutations detected in HSCs were not present in BM cells, whereas we could identify known aCH mutations in BM cells. Our data reveal that the mutational and clonal landscape in aged HSCs is uncoupled from aCH.

## Results

To reduce false-positive SNVs in single-cell sequencing, we devised a simple yet robust approach to variant identification using HSCs sorted from murine and human samples. In the first step, we performed single-cell RNA sequencing (scRNA-seq, RNA-based, SMARTseq technology) and single-cell assays for transposase-accessible chromatin (scATAC-seq, DNA-based) on paired daughter cells from the first *ex vivo* division of individually sorted HSCs (Lin^-^ Kit^+^ Sca-1^+^ Flk2^-^ CD34-cells) from young (8–16-week-old C57BL/6 mice) and old (> 80-week-old C57BL6) mice (**Fig. 1a** and **Fig. S1a, left panel**). After variant calling on individual cells, we filtered for position-matched SNVs in pairs of daughter cells and scored only those variants present in both daughter cells (joint filtering) as true variants and included them in downstream analyses (**Fig. 1b**). This critical filtering step disqualified about 60-80% of the raw SNVs (**Fig. 1c** and **Fig. S1a**, and **Table S1**). To assess the rate of false-positive events in our approach, we applied a Monte Carlo simulation by performing chromosome-controlled, positional random reshuffling of mutational events in each single cell and counting matches across daughter pairs. In 1000 iterations for each daughter pair, the overall mean random positional match between daughter pairs was less than one (scRNA-seq = 0.028, **Fig. 1c** and **Table S1**; scATAC-seq = 0.0047, **Fig. S1b**, and **Table S1**). Because this first simulation step did not account for the SNV match, we adjusted the results to reflect the probability of independent random base-substitution matches per position (P = ¼ x ¼ = 0.0625). The mean final random positional and mutational match between daughter cells was close to zero (scRNA-seq = 0.0018, scATAC-seq = 0.00029; **Fig. 1c** and **Table S3**), which was significantly lower than the empirical SNV count (p-value < 0.001) (**Fig. 1c** and **Table S3).** Therefore, the co-occurrence of an identical SNV in two daughter cells is a non-random event and represents a pre-existing variation in the mother HSC.

**Figure 1.**
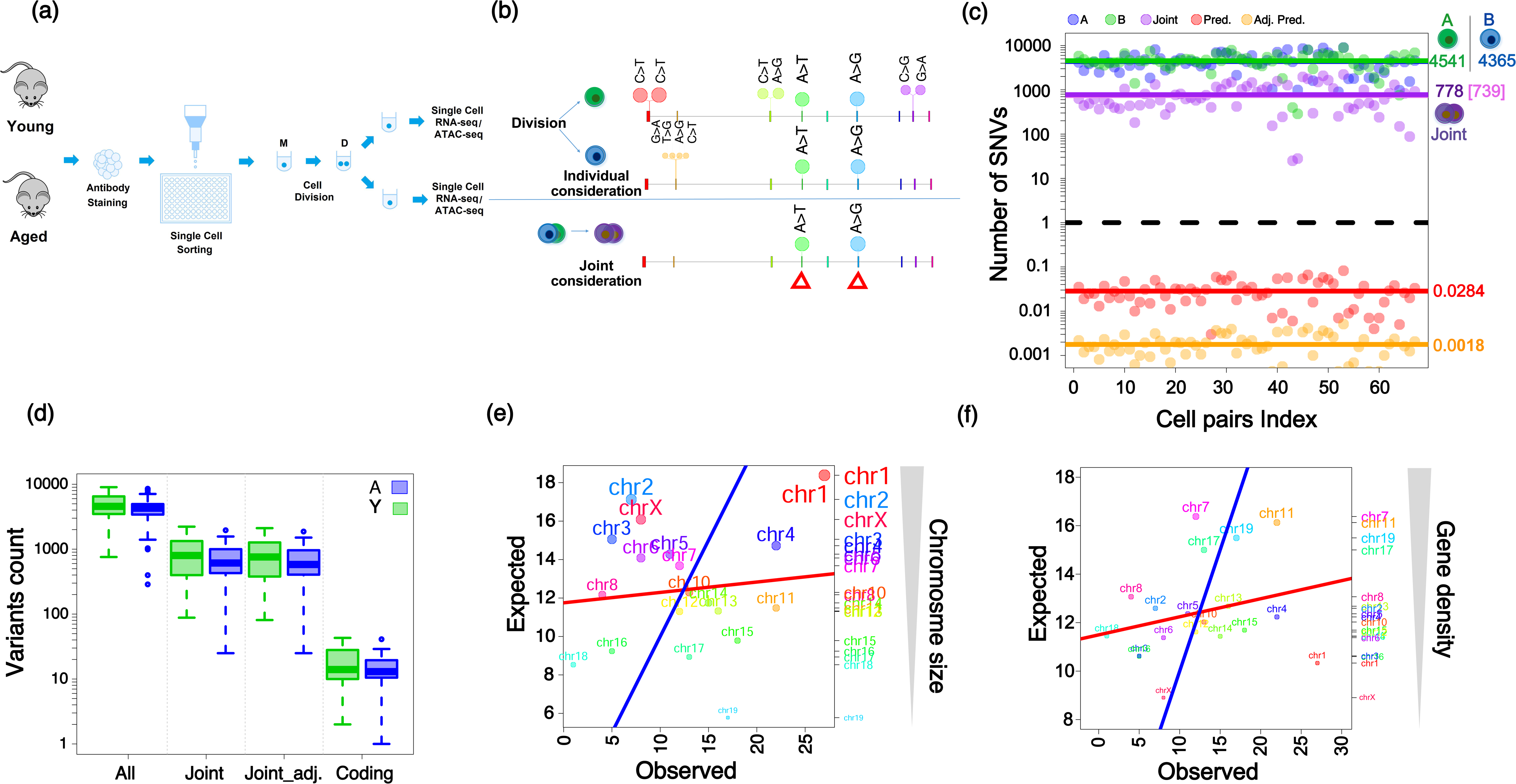
**(a)** Schematic representation of the experimental design. Long-term HSCs (LT-HSCs; Lin-Sca1+Kit-, CD34-, Flk2-) were sorted from young and aged mice. Every single cell (M) underwent one round of cell division to give rise to two daughter cells (D). Each daughter cell was placed in a separate well and subsequently subjected to single cell RNA sequencing. **(b)** A depiction of the variant filtering rule used in the SNV analysis. We first performed SNV calling for each cell separately (individual consideration = green and blue cells; lolliplot shows individual SNVs). We then used a joint variant filtering approach where we kept those variants that occur in both daughter cells (Simultaneous consideration = depicted by overlapping purple cells and red boxes overlaid on the lolliplot highlighting the SNVs fulfilling the criterion). **(c)** Variant count in each cell pair before and after applying our filtering rule. Green and blue points represent SNV counts in each daughter cell (A=green, mean (𝑥̅) = 4541; B=blue, mean (𝑥̅) = 4365; represented by colour matched lines). Purple points indicate the number of position matched variants in daughter pairs (Joint=purple, mean (𝑥̅) = 778). The final mean count of position and base substitution matched SNVs is 739. Dashed line indicates mean variant count value of 1. Red points represent the predicted number of position matches in a Monte Carlo Simulation (MCS) analysis based on position reshuffled daughter pairs (mean (𝑥̅) = 0.0284; 1000 iterations per daughter pair). Similarly, the orange points and line represent the MCS results after adjusting for the probability of base substitution match for each positional match (mean (𝑥̅) = 0.0018). **(d)** Distribution of variants per cell in young and aged HSCs based on the scRNA-seq (Young = green box; Aged = blue box). Shown here are all variants before filtering (All), variant count after filtering for positional matches (Joint), positional and substitution matches (Joint_adj), and finally SNVs that strictly fall in exons (Coding). It can be noted that young and aged HSCs show no difference in variants counts across all categories, albeit a subtle reduction in Aged HSCs. **(e)** Coding variant distribution by chromosome size in the scRNA-seq dataset. The observed number of variants per chromosome (x-axis) plotted against the expected count relative to the chromosome size. Red line represents fitted line showing the relationship between expected and observed variant frequency while blue line depicts the y=x of the expected frequency i.e. alignment of the red and blue lines would indicate chromosome size dependence of variant counts. Using a goodness-of-fit test, we observed that this is not the case and variant count frequency is independent of chromosome size (𝜒^2^ = 41.89, p-value = 0.0011). Second y-axis (and grey-filled arrow) shows a descending order of chromosomes sizes. **(f)** Coding variant distribution by chromosome gene density (number of genes per million bases, genes/Mb) in the scRNA-seq dataset. Similar to figure 1e, the observed number of variants per chromosome (x-axis) plotted against the expected count relative to the gene density. We also observed here that variant counts are independent of gene density in a given chromosome (𝜒^2^ = 69.38 p-value < 0.001). Second y-axis (and grey-filled arrow) shows a descending order of chromosomal gene density.

The mean number of position-matched variants per HSC was 778 and 587 for murine HSCs in the scRNA-seq and scATAC-seq experiments, respectively. Remarkably, the frequency of SNVs in the scRNA-seq and scATAC-seq, identified using independent experimental protocols and analyses, fell within a very similar range of counts in HSCs. The fact that both ATAC-seq and RNA-seq show a very similar range of mutations implies a likely moderate influence of such factors on the overall call rate of SNVs in HSCs. Considering SNVs found in daughter cells will thus allow for a reliable, reproducible, and quantitative determination of true SNVs in single cells without the need for any further statistical or context-dependent mathematical correction to select for true SNVs.

In the next step, we slightly modified the approach to identify true SNVs in conditions where daughter-pair information is either experimentally unavailable or not feasible. In this approach, still based on the paired daughter scRNA-seq dataset, we scored an SNV as true when it was present in at least two single cells. When comparing this new set of SNVs to the daughter-pair-matched SNVs, we detected all of the previous SNVs, and the total increased by only an additional 10% of all SNVs called across all cells in the single-cell murine RNA-seq dataset. This increment could be attributed to the inclusion of heterozygous SNVs, in addition to the strictly homozygous variants detected via daughter-matched analysis, that might be linked via a two-generation relationship. Filtering for 2 identical SNVs in a dataset of single HSCs provides a list of true SNVs in individual HSCs with an extremely high level of accuracy, while the method likely still underestimates the total number of SNVs present in single HSCs. The absolute number of SNVs in HSCs identified with our approach might thus still need to be considered very carefully, as it will not call heterozygous mutations in individual cells that are not present in any other cell. Furthermore, mutations in alleles that are not transcribed in the cell of interest will also not be scored. The strong advantage of this simple filtering approach is that it identifies real SNVs in single cells without any additional modeling, allowing for in-depth quantitative comparisons of SNV frequencies among single HSCs and populations.

Interestingly, by comparing the number of SNVs between young and aged HSCs in paired-daughter RNA-seq and ATAC-seq datasets, our approach revealed no difference in the number of SNVs per cell between the two age groups, and, if anything, a subtle reduction in the total number of SNVs in aged HSCs (**Fig. 1d**, **Fig. S1d,** and **Table S2**). This pattern holds for both scRNA-seq and scATAC-seq datasets and is also observed for SNVs strictly in exonic regions (considering only coding SNVs) (**Fig. 1d**, **Fig. S1d,** and **Table S2**). A goodness-of-fit test showed that the distribution of SNVs in both scRNA-seq and scATAC-seq datasets is independent of either chromosome size or gene density (**Fig. 1e-f** and **Fig. S1d-e**). The data thus demonstrate, contrary to a widely accepted paradigm, that the number of SNVs per single HSC is not increased in aged compared to young murine HSCs. The moderate but significant increase in overall mutation frequency in aged hematopoietic progenitor cells previously reported by other approaches (Moehrle et al., 2015) is thus likely independent of the mutational load of HSCs.

We next examined the nature of the SNVs in more detail to gain insight into their genic context. Of the 236 total coding SNVs, 88 were shared between the two age groups, while 70 and 78 were unique to young and aged HSCs, respectively (**Fig. S2a**). At the gene level, 38 genes were unique to young HSCs and 31 were specific to aged HSCs, while 51 were shared (**Fig. S2a**). Interestingly, neither the 38 young HSC-specific nor the 51 shared SNVs between young and aged yielded significant Gene Ontology (GO) terms in a GO enrichment analysis. Notably, epigenetic regulation of gene expression, chromatin organization, and metabolic processes were statistically significantly enriched based on the 31 aged HSC-specific genes (**Fig. S2d**).

We further focused on aged HSC-specific variants to identify both qualitative and quantitative mutational dynamics in HSCs during aging. Among the 31 genes unique to aged HSCs, the median number of distinct SNVs per gene was one, and the median per cell was two (**Fig. S2b**). *XLR4c* was most frequently marked by an SNV in aged HSCs. *XLR4c*, whose function is still unknown, is an imprinted X-linked gene expressed independently of X-chromosome inactivation in immature hematopoietic cells (Grigoryan et al., 2021) and in differentiated T-cells (Raefski and O’Neill, 2005). Interestingly, we did not find in aged (or young) murine HSCs SNVs enriched in genes linked to hematopoietic disease (leukemia) in either mice or humans.

To rule out expression bias as an underlying reason for age group-specific variants, we tested whether the expression levels of genes harboring SNVs in young or aged HSCs show differences between the two age groups. We first examined the corresponding log mean Fragment Per Kilobase Million (FPKM) expression of 109 of the total 120 genes (all coding SNVs/genes considered) and 30 of 31 (aged HSCs-specific SNVs/genes) whose gene-level expression across some or all cells in either the young or aged group was quantified. We observed that these genes did not show a statistically significant difference in expression between the two age groups (**Fig. S2c**). SNVs, therefore, in general, did not influence the level of gene expression. In a two-group, gene expression-based, unsupervised hierarchical clustering, we checked whether young or aged HSCs showed a significant odds ratio of cluster membership. This analysis is important because it tests whether the expression levels of these genes tend to group cells by age group. In line with our test of the significance of the difference in expression, Fisher’s exact test of cluster membership showed no statistically significant difference, confirming that there is no directional trend in gene expression of our selected genes specific to an age group (Tables **S4**, **S5**, and **S6**). This finding rules out the possibility that our ability to detect these SNVs specifically in the aged samples might be simply linked to a distinct level of expression.

While aging-related clonal evolution and clonality in hematopoiesis are widely discussed in humans, almost nothing is known about these events in mice. To determine clonal relationships among the HSCs analyzed, we next evaluated cell-to-cell similarity based on the identified coding SNVs. This approach quantifies the genealogical relationship among HSCs and serves as a surrogate for clonal relationship and thus clonality among HSCs. To this end, we performed consensus clustering using a pairwise Hamming distance between cells within each age group. In young HSCs, we observed one major cluster, with 73% of cells assigned, while the remaining cells formed singleton branches (**Fig. 2a**). Aged HSCs, in contrast, formed four distinct clusters, with 43%, 26%, 11%, and 9% of cells in each cluster, respectively, for a cumulative total of 89% (**Fig. 2a**). The remaining cells formed singleton branches. Aged HSCs thus showed a distinct and structured subgrouping, strongly implying that aged HSCs are distinct clones that have been selected over time. Young HSCs generally showed high cell similarity (lower distance) and no subclustering.

**Figure 2.**
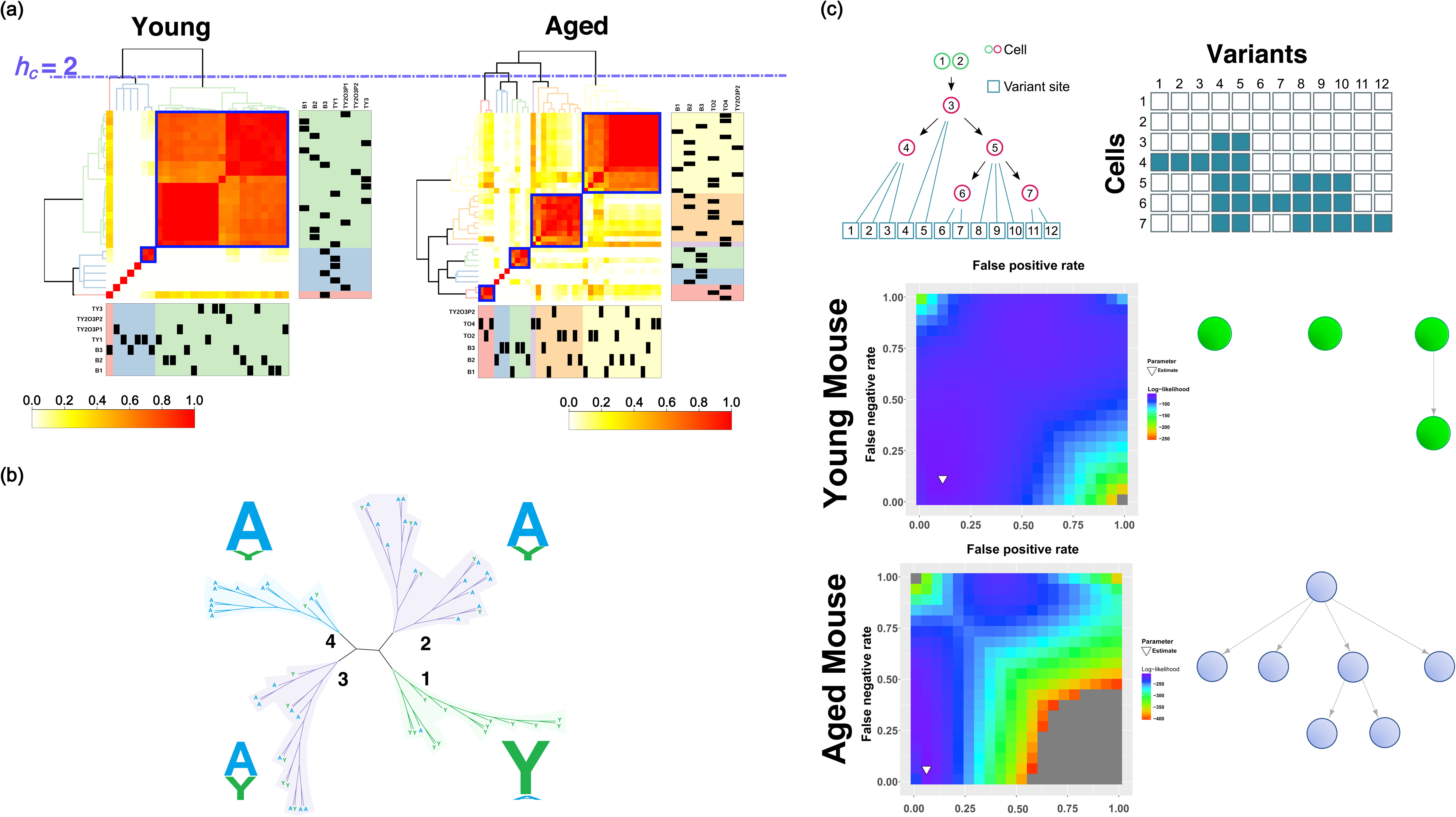
**(a)** left panel: Hamming distance-based consensus clustering of young HSCs. Mouse numbers are indicated below and on the right side of the heatmap. At a cluster height of 2, we observed 1 major cluster (light green) where more than 70% of the cells were grouped into. Both the light blue and light red clusters are single cells except one sub-cluster with 2 members. Right panel: Heatmap based on consensus clustering of aged HSCs revealed 4 main clusters at a cluster height of 2 (yellow, pink, light green, and red), with 43%, 26%, 11%, and 9% of cells being assigned to them. Light blue and purple clusters contain a single member. **(b)** Neighbor-Joining (NJ) Phylogenetic tree based on the pairwise Hamming distance between all cells irrespective of age group. Four major branches have been labeled as either A (Aged; blue; three branches) or Y (Young; green) based on the majority of cells in each branch. Size of labels (A or Y) is proportional to the number of cells representing the age group for a given branch (see main text for details), the dominant age category being drawn first. **(c)** Coding SNVs based evolutionary relationship among cells (cellular/clonal evolution) in a given mouse using the scRNA-seq dataset (here are shown mouse ID B2 in young and aged groups). The tree building model has been adapted and modified from Ross and Markowetz, 2016). Briefly, the variants table (left part of the model) is used to hierarchical order and assign positions of cells in the tree, with incremental mutations on preexisting ones leading to the placement of a cell lower in the hierarchy (Cells at the apex are founder cells relative to the pool of cells being analyzed). Cells are thus placed in the hierarchy based on maximum likelihood-based estimation based on a nested structure of the mutational pattern. Cells not linked with arrows indicate that there is no sufficient deducible relationship based on the provided SNV data. Here are shown representative plots of one young and one aged mouse. It can be noted, for the mice presented here, that aged HSCs show a simple proliferation like a hierarchical model while the young HSC show little or no intercellular evolutionary relationship. The heatmap show profile of false positive rate (FPR, x-axis) and false negative rate (FNR, y-axis) in a given analysis, the inverted triangle within the plots indicating the estimated FPR/FNR for that analysis. See supplementary figure 3 for plots of all mice in the study.

We then combined the young and aged HSCs data and calculated cell-to-cell pairwise Hamming distances irrespective of group membership. Based on this, we constructed a Neighbor-Joining (NJ) phylogenetic tree that diverged into four major branches. Branch 1 contained predominantly young HSCs (13/14 or 93% of members), with only one aged HSC cell pair assigned to it (**Fig. 2b**). The second and third branches contained mainly aged HSCs (75% and 60%), while the last branch had 10/12 or 83% aged HSCs (**Fig. 2b**). Branches were defined as either young or aged based on a chi-square test of the null hypothesis of proportional assignment of cells from both age groups. Accordingly, three branches were defined as “Aged” and one branch was defined as “Young” (p-value < 0.05; **Fig. 2b**). The data, therefore, imply that aged HSCs, assigned to divergent genealogical clusters or branches, have evolved to form distinct genetic sub-entities, whereas young HSCs are predominantly grouped into one large branch with little sub-clustering. These findings are in line with the age group-based clustering analysis (**Fig. 2a**). In a final step, we performed a clonal lineage tree analysis of HSCs from individual mice to obtain insight into the genetic and genealogical relationships among HSCs from individual animals and to construct clonal hierarchy models using an established maximum likelihood approach that deciphers the clonal tree by evaluating the order and overlap of mutational events observed in the cells (Ross and Markowetz, 2016) (an adapted and modified model is shown in **Fig. 2c**, top panel). We generated clonal trees for all 12 mice (6 Young; 6 Aged; **Table S8 and S9**). Eleven of the 12 mice had false-positive and false-negative rates at or below 0.25 (**Fig. 2c** and **Fig. S7**). Interestingly, we observed higher and more ordered cell-to-cell relationships among HSCs from individual aged animals that fit a simple clonal proliferation-like model, compared with HSCs from individual young animals (**Fig. 2c** and **Fig. S7**), findings that are further consistent with the distinct grouping we observed in our consensus clustering and phylogeny analyses in **Figs. 2a** and **2b**. Overall, the data thus imply that HSCs in aged mice, compared with HSCs in young mice, stem from only a few founder clones, indicating that clonality exists among aged stem cells.

Whether clonality begins within the HSC pool during aging in humans and whether this is linked to heterogeneity among HSCs remain matters of debate. Identifying SNV dynamics in individual human HSCs is therefore central to our understanding of the role of HSCs in aCH.

To determine the frequency and type of SNVs in individual human HSCs, we generated and analyzed human single-cell RNA-seq datasets from HSCs (Lin-CD34+^SSc^ ^low^ CD38-CD90+CD45ra-; **Fig. S1a, right panel**) isolated from BM obtained from 4 young (<30 years) and 4 aged (>65 years) healthy individuals, with 128 single cells sequenced per donor. Cells were collected individually without daughter cell pair information. High-confidence SNVs, as discussed in previous sections, were scored when present in at least two single cells. To assess the false-positive rate for this dataset, we again performed a Monte Carlo simulation by permuting two cells at a time, reshuffling the detected SNVs, and checking for random matches. Of the average ∼8000 SNVs per cell, we observed that the average number of true SNV matches per two cells was ∼100, while the median and average of random matches per 1000 iterations of a pair of cells were 0 and ∼0.01, respectively, with a maximum of 4-5 positional matches and 0.3 base substitution matches (**Fig. 3a, top left panel**). In conclusion, the co-occurrence of an identical SNV in two stem cells is rarely a random event, and, as a consequence, SNVs fulfilling this criterion are “high-confidence” SNVs.

**Figure 3.**
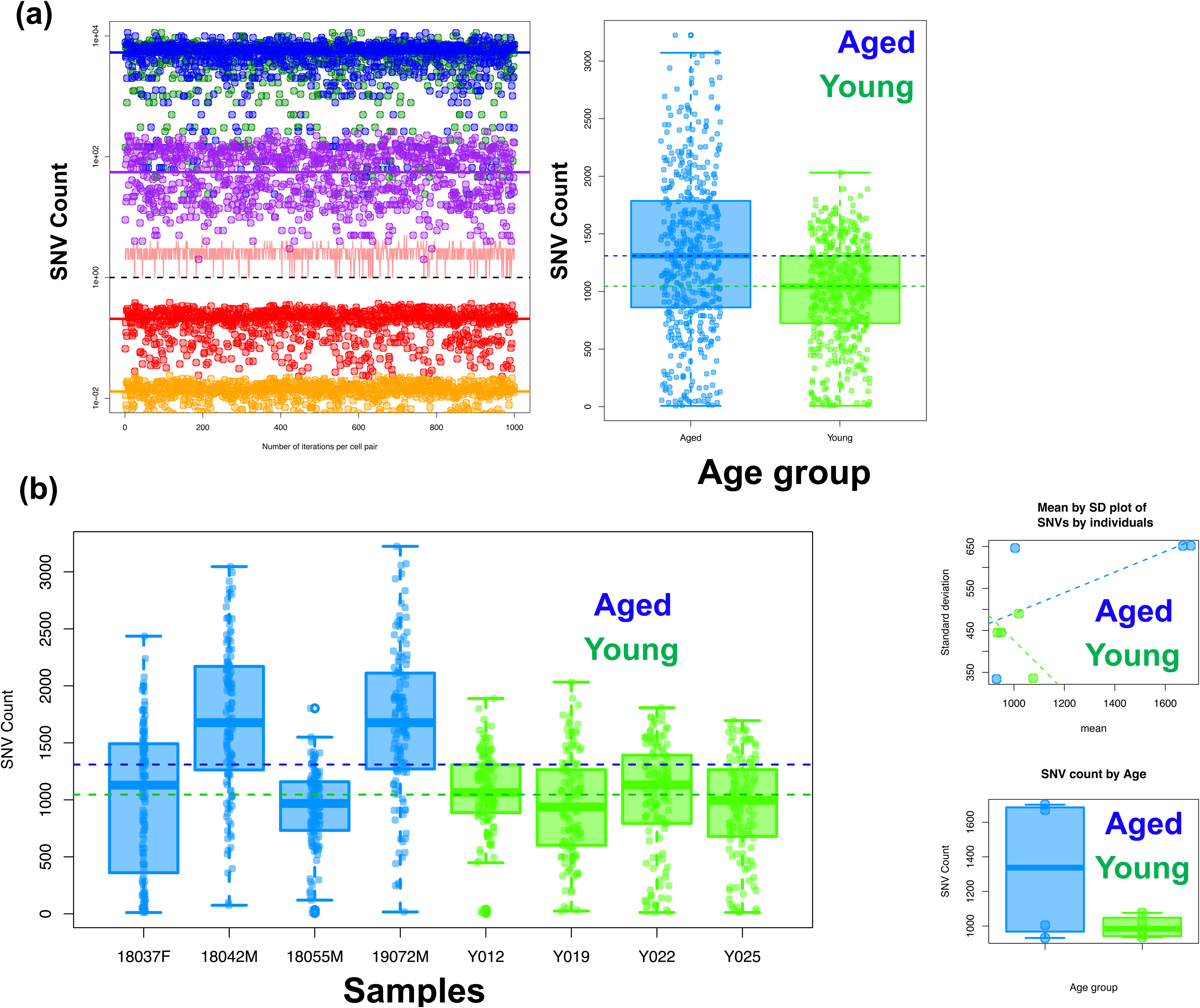
**(a).** Comparative profile of various filters as applied to the raw SNV counts in the mouse single-cell RNA-seq dataset. By setting all SNVs to a 100 % count (blue area), we show that our daughter pair matched SNV count accounts for 3.37% of the called SNVs (red). The modified approach of considering an SNV if it is present in at least two cells are 13.45% (green) of the total SNV count. **(b). (top left panel)** A Monte Carlo simulation analysis of a representative sample of the human single-cell RNA-seq dataset. Blue and green points represent number of SNVs in two randomly picked cells per iteration (median ∼ 8000 SNVs per cell; blue and green line). Purple points depict the number of matched SNVs between the picked cells (median of ∼80 SNVs per cell pair). Red points represent the number of randomly matched SNVs between those cells after the SNVs in each cell were reshuffled and positionally matched (red line shows a median match of < 1 positional SNV matches; maximum number of matches per iteration in the random analysis is shown by the pink line). The orange points show the number of SNVs when we adjusted the positional match frequency to account for the probability of random base match per position (see text for details). **(top right panel)** An overall comparison between young (green) and aged (blue) SNV counts using the approach of at least two cells per SNV in a given sample. We could show a slightly higher (∼1.25 times median increment) SNV count in aged samples as compared to the young. **(bottom left panel)** Individual samples SNV count profiles of young and aged samples. **(bottom right panel)** We noted higher consistency in median in SNV count in young samples as compared to aged ones as shown here by mean by standard deviation (sd) plot (note the higher sd in aged samples) and plot of the median SNVs by sample across both age groups (one can see the higher interquartile range in aged samples).

To test the hypothesis of increased DNA mutation load in human HSCs with aging, we compared the number of SNVs per cell between young and aged HSCs. We observed a slightly higher mutational frequency per cell in aged human HSCs compared with young HSCs (**Fig. 3a, top right panel)**. Additionally, we noted greater variability in SNV frequency in aged individuals compared with young ones (**Fig. 3b, bottom panels)**, consistent with a general increase in variability and heterogeneity among identical cells with age (Bahar et al., 2006). A direct comparison of the mean SNV count across cells between young and aged individuals was not statistically significant, indicating that young and aged HSCs have a similar mutational load, in line with our findings in the murine single-cell analysis (**Fig. 3b, bottom panels)**. When we performed an age group-specific overlap of the identified SNVs between individuals, we observed that 7371 and 5872 SNVs (corresponding to 737 and 597 genes) were shared between aged and young individuals, respectively (**Fig. S4**). By taking these overlaps, we also showed that 3779 SNVs (or 493 genes) were common between young and aged samples. These common SNVs and genes were numerically larger than the age group-specific SNVs. Although we still observed a slightly higher number of SNVs in aged samples compared with young ones, these findings support that aged stem cells largely maintain the young mutational landscape and that therefore an aged microenvironment might be more crucial in defining clonal selection.

Due to the high number of single cells per individual donor in the human dataset, we were able to perform in-depth, individual-based clonality analyses. Interestingly, we did not observe significant differences in clone numbers between young and aged individuals (**Fig. S5**). To assess whether the clones we identified in young and aged samples revealed transcriptional programs reminiscent of an aging phenotype, we first employed a deep learning algorithm to generate a high-confidence aging signature that could accurately predict and classify HSCs as young or aged. The deep learning analysis confirmed prediction accuracy using a validation subset of single cells not used during model training (∼93% accuracy; p-value 1.88-e66; **Fig. S3**). The top 1% of predictor genes with the highest correlation coefficient to the deep learning model showed statistically significant enrichment for HSC stemness and aging signatures (**Fig. S3)**. To our knowledge, this is the first validated single-cell-based aging signature for human HSCs.

Pairwise differential expression analyses allowed us to re-group HSC into two major clones based on their expression profiles, with one group containing a branching-out clone and the remaining clones falling into the second group (**Fig. 4a, left** and **middle panel**s show a representative sample). A GSEA of the two major clones, specifically in aged samples, indicated that the branching-out clone shows upregulation of an aging signature and downregulation of the HSC signature (**Fig. 4a, right panel**). This supports the idea that, in an aged individual, not all cells and clones age at the same rate, and only a fraction of cells/clones acquire expression changes linked to aging.

**Figure 4.**
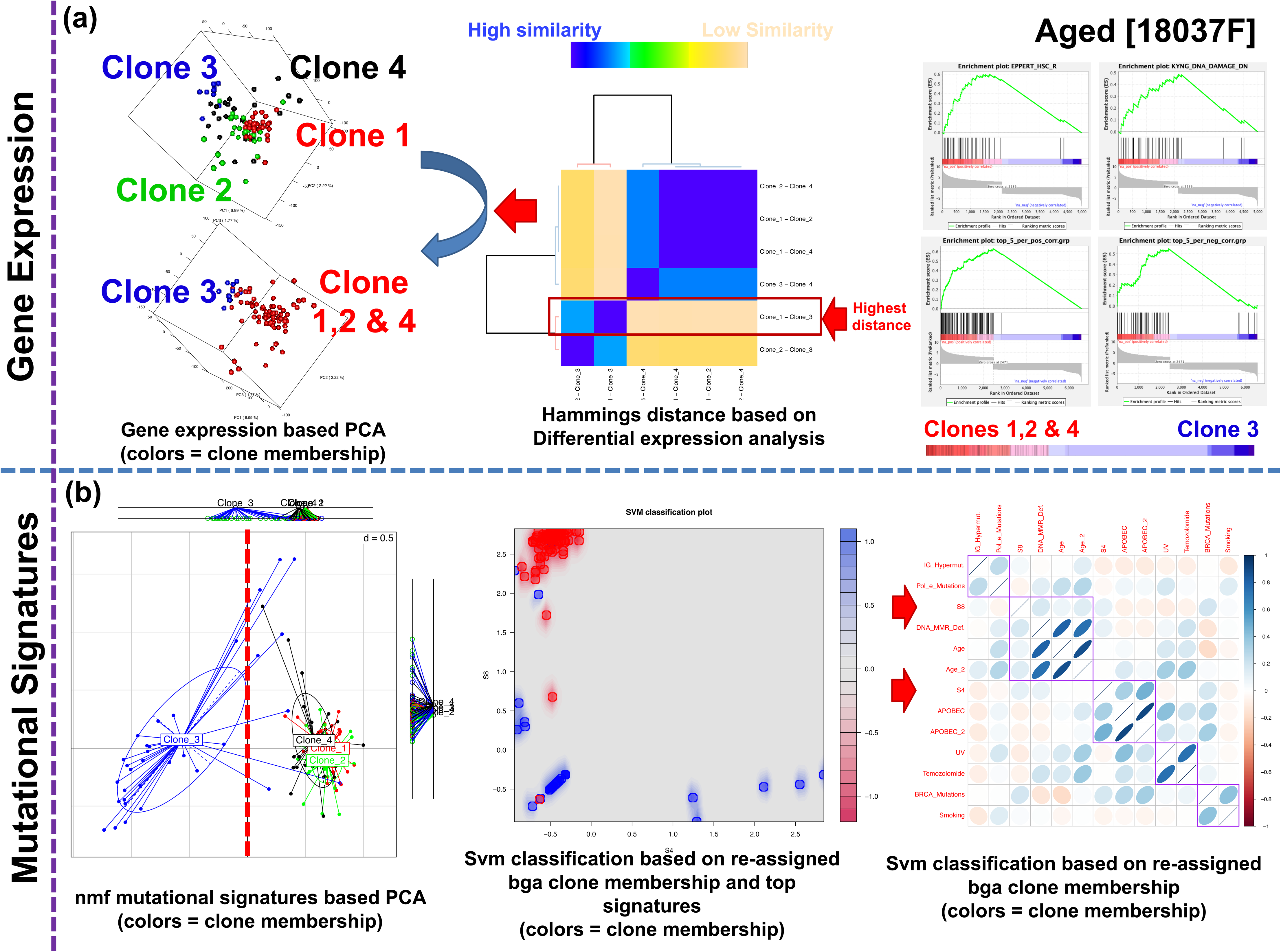
**(a).** A representative aged sample clonality and clonal grouping-based gene expression analysis. We detected four clones in this sample (see methods for details on clonality analysis). A pairwise differential expression followed by Hamming’s distance between the pairs of comparisons reveals the most distant set is clone 1 to clone 3. The rest of the pairs had high similarity (blue part of heatmap), except another pair of comparison including clone 3. A 3D principal component analysis (PCA) using original group coloring (top part; 4 colors) and a recolored one based on the pairwise differential expression analysis (lower part; 2 colors where clone 3 is colored blue while the other clones are colored red). We also performed GSEA (right panel) where we show hematopoietic stem cell and DNA damage signature show significant upregulation in the clones 1,2, and 4 as compared to clone 3. We also performed GSEA of the top 5% predictors from the deep learning analysis and could show that they show upregulation in the clones 1,2, and 4, indicating that clone 3 is not only genetically but also transcriptional more distinct from the other clones. **(b).** A non-negative matrix factorization (nmf) mutational signatures (see methods for details) matrix based between groups analysis (bga) shows that clone 3 is more distinct from the other clones **(left panel)**, in line with what we showed with the transcriptional analysis. We then tested 10 signatures in a pairwise fashion (using top scoring pairs and support vector machines, svm) to identify the pairs with the highest discrimination score. Here is shown the svm plot **(middle panel)** of signatures 4 and 8, where clone 3 in blue shows clear segregation the rest of the clones (red). **(right panel)** a correlation between signatures 4 and 8 and publicly available signatures with known biological association. We could show that S8 shows positive correlation to aging and DNA mismatch repair deficiency signatures while S4 is positively correlated to APOBEC signatures.

Recent work showed that a mutation and its 5’ and 3’ flanking bases generate specific motifs whose frequencies can be mathematically analyzed to deduce mutational signatures that inform the likely origin of the mutation (Alexandrov et al., 2013a; Alexandrov et al., 2013b; Alexandrov and Stratton, 2014; Nik-Zainal et al., 2012). These mutational signatures primarily grouped the clones into two major subclones. Notably, the clone that was distinct in the gene expression analysis also formed a separate cluster along principal component 1 (**Figure 3b, left panel**). In the second step, top-scoring pair and support vector machine (svm) analyses based on the top signatures (**Fig. 4b, middle panel**; **Fig. S8; Fig. S9**) were correlated with published mutational signatures. The aged HSC somatic mutational signatures showed a significant correlation with aging and DNA mismatch repair (MMR) deficiency signatures, whereas young HSC-based mutation signatures did not show any significant correlation with any of the published signatures (**Fig. 4b, right panel**; **Fig. S9**). Similarly, in a Principal Component Analysis (PCA) of the murine dataset, 40 mutational signatures were generated that explained more than 99% of the variance in the dataset (**Fig. S6a**). We analyzed the signatures using a top-scoring pair (tsp) analysis and identified a pair of signatures with the highest discrimination score in classifying young and aged HSC (S5-S13 pair; **Fig. S6b**). A cross-validation Odds-Ratio (OR) analysis of all signature pairs confirmed that the S5-S13 signature pair-based group assignment was statistically highly significant (p-value 0.0042; tsp/OR significant signatures overlap is shown in **Fig. S6c**). Using a support vector machine (svm), we detected a higher sub-clustering tendency in aged HSCs (5 centroids) compared with young HSCs (only 2 centroids) (**Fig. S6e**). This suggests that a less diverse aged HSC pool evolves from a heterogeneous pool of highly individualized young HSCs (and thus only 2 distinct clusters could be detected). We then generated 40 nmf-based signatures and performed tsp analysis. We found two pairs of signatures with the highest young-aged HSC discrimination score (S12-S26 and S12-S35; **Fig. S6c-d**). In the final step, similar to the human dataset, we correlated the three signatures (S12, S26, and S35) with published signatures with known biological associations. We observed that S26 and S35 showed a highly significant correlation with aging and DNA mismatch repair (MMR) signatures (**Fig. S6d**). The probable underlying cause of the aging-related mutation signature has been linked to elevated rates of spontaneous hydrolytic deamination of 5-methylcytosine, which results in C>T transitions (Alexandrov et al., 2013a), while the MMR signature has been described as characteristic of malignancies with defective DNA mismatch repair (Boland and Goel, 2010). Therefore, SNVs in aged HSCs (aka HSC aging signature) seem to stem from a basic chemical process (deamination) (Lewis et al., 2016) and from errors in the repair mechanism (MMR) that resolves mismatches resulting from deamination (Jones et al., 1987). Interestingly, defects in MMR have been previously implicated in the aging of human hematopoietic progenitor cells (Kenyon et al., 2012).

aCH is generally defined by a set of frequently mutated genes. SNVs in these genes are therefore considered a hallmark of clonal hematopoiesis in the elderly (Beerman et al., 2014; Rossi et al., 2007). We examined 11 genes most frequently associated with aCH in our SNV dataset, including *TET2*, *JAK2*, *DNMT3A,* and *BCOR1* (Buisman and de Haan, 2019; Jaiswal and Ebert, 2019; Steensma et al., 2015). None of these genes were differentially expressed between young and aged HSCs, which excludes any skewing in SNV calling due to differential expression of these genes (**Fig. S10,** upper panel). We found SNVs in 9 of the 11 genes tested, with *JAK2* and *TET2* showing the highest recurrence across individuals. HSCs from 7 of 8 individuals (young or aged) harbored these mutations, with an overall frequency of less than 10% among all HSCs analyzed per individual for most mutations, except a novel *JAK2* mutation with a frequency of 15% (**Fig. 5a** and **Fig. S10,** lower panel). The previously unreported *JAK2* SNV (gene position A2741T; protein position K914M) showed the highest recurrence among all 11 genes tested across all HSCs and individuals. Interestingly, the *JAK2* SNV was among the subset of SNVs identified as critical in our clonality analysis, implying that it contributed to shaping the observed HSC clonal landscape. All other SNVs (**Fig. S10,** lower panel) were not identified as significant contributors in our clonality analysis. Furthermore, a gene expression-based GSEA comparing cells grouped by *JAK2* SNV status among HSCs revealed that cells with the *JAK2* SNV show downregulation of the HSC signature and upregulation of aging signatures, making them comparatively even more “aged,” consistent with reduced fitness of murine HSCs that lack *Jak2* (Kent et al., 2013). Do mutations in *JAK2* arise in a manner different from the other SNVs detected in human HSCs (**Figure 4b**), which might then explain their relatively high frequency? To address this question, we identified the SNV mutational signatures on a representative sample, using the JAK2 mutation as a grouping parameter. The two signatures with the highest discrimination score between *JAK2*-mutated and unmutated cells revealed that one of the signatures still showed a significant correlation with aging and DNA MMR, in addition to UV and Temozolomide signatures (**Fig. 5b, right panel**), similar to the bulk of aged human HSCs. SNVs in JAK2 are therefore likely not generated by mechanisms distinct from those that generated the other SNVs found in aged HSCs.

**Figure 5.**
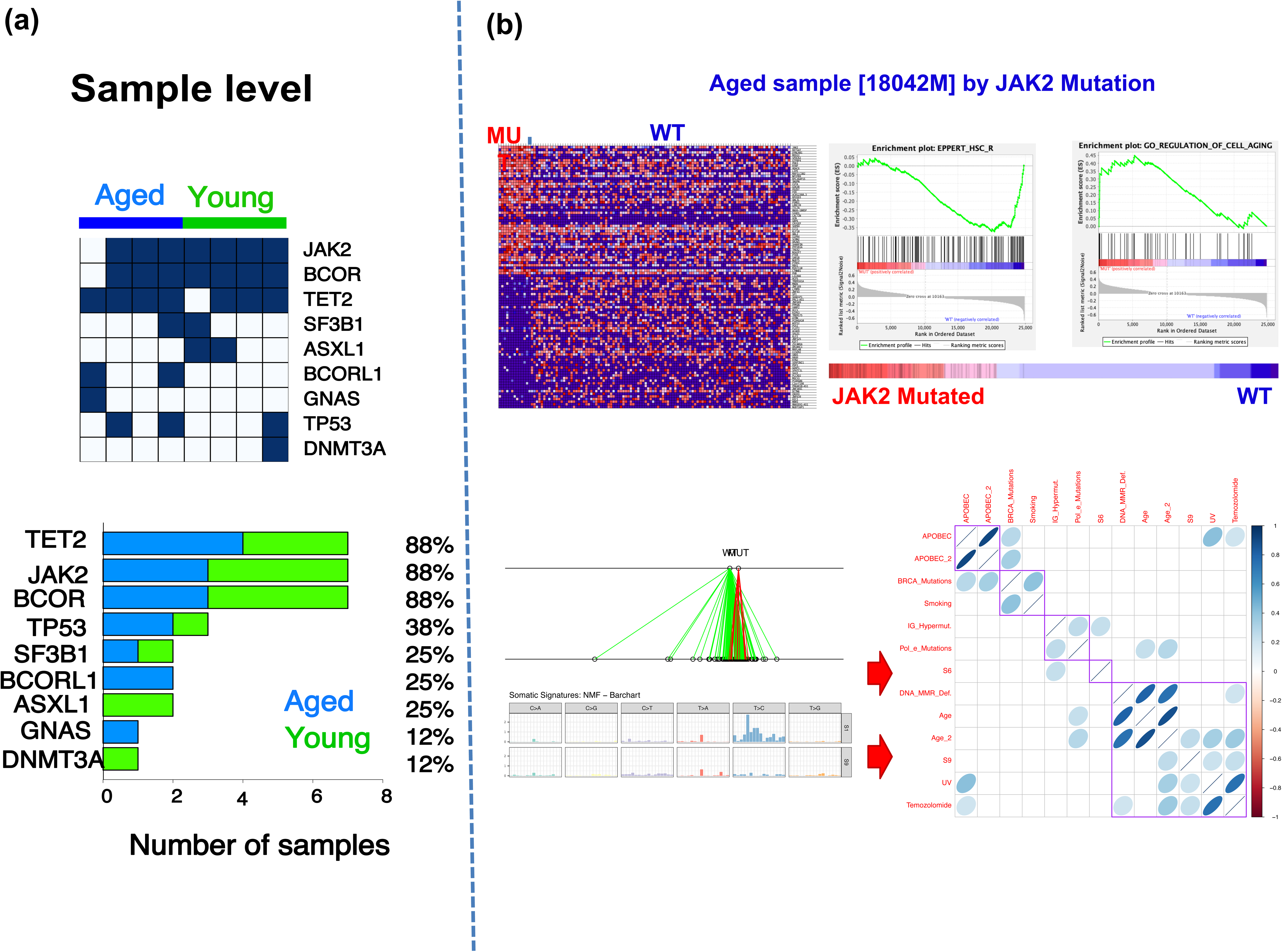
11 mutations reported to be relevant to clonal expansion in hematological malignancies including *TET2*, *JAK2*, *DNMT3A* and *BCOR1* were looked at in our human single-cell variants dataset. **(a)** Distribution of detected mutations in genes across samples. Mutations in *JAK2*, *TET2*, and *BCOR* were detected in 7 out of 8 individuals while only one individual harbored a *DNMT3A* mutation. **(b)** Since the *JAK2* mutation was a unique and had relatively high recurrence across cells and individuals, we grouped cells as JAK2 mutated (MU) and non-mutated (WT) and performed GSEA on a representative aging sample (upper panel). We observed that cells with mutations show downregulation of stem cell signature and upregulation of genes involved in regulation of cell aging. **(lower panel)** We identified two mutational signatures with the highest discrimination score between *JAK2* mutated and WT groups (S6 and S9) and S9 showed positive correlation to mutational signatures associated with ultraviolet (UV) and Temozolomide.

We next performed targeted DNA sequencing of bone marrow (BM) cells from the samples from which HSCs were isolated to assess whether SNVs identified in HSCs were also present in the bulk BM. Coverage of the target region ranged from 250 to 3500, with a median of ∼1500 (**Fig. S11**). The minimum allelic frequency of the mutations we detected in the bone marrow was 0.149 (**Fig. S12,** lower middle panel). As anticipated, we detected known aCH mutations in BM cells, but none of the SNVs identified in HSCs were detected in BM (**Fig. S12**). Additionally, the ranking of genes by sample recurrence was distinct. For instance, *ASXL1* was the most recurrently mutated gene in BM cells, whereas *JAK2* and *TET2* were mutated in HSCs (**Fig. S12** and **Fig. 5**).

To validate our observations in this study that mutations in HSC are generally low and do not contribute to the larger pool of SNVs in the bone marrow (the bone marrow mutational profile largely reflects more abundant cells with higher mutational load), we performed mutational analysis of single-cell data from stem and progenitor cells using the recently published dataset (Bandyopadhyay *et al.,* 2024). Although 10x genomics-based single-cell RNA-seq datasets are generally not suitable for mutational analysis, they are sufficient for questions that primarily examine the comparative frequency and similarity of SNVs across cell types. The cells we analyzed were CLP, Cycling HSPC, GMP, HSC, MEP, and MPP. In a randomly selected subset of reads, we performed all SNV-calling and filtering steps discussed in this study (see the methods section). We first examined the overall SNV count. As shown in **Fig. S12a**, the lowest median SNV count was in HSC, followed by CLP. To assess the degree of similarity among specific SNVs, we built a hierarchical clustering based on Hamming distance (heatmap; **Fig. S12b**). GMPs and MEPs showed the highest similarity, while the other cell types formed distinct arms in the dendrogram. Furthermore, a between-group PCA revealed that MPPs are distinct from other cell types along PC1. Although HSC formed a distinct cluster in the two-dimensional plot, PC2 showed overlap between MPP and HSC, while GMP and Cycling HSPC (on the positive PC2 axis) and MEPs and CLPs (on the negative PC2 axis) showed strong overlap (**Fig. S12c**). In the final step of the analysis, we examined the proportion of SNVs shared between cell types (pairwise). Although cells shared a fraction of their SNVs, cell-type-specific SNVs accounted for a large proportion of the total (**Fig. S12d**). SNVs shared between HSC and other cell types (red ribbons) contributed to a small fraction of each cell type. These results indicate that not all mutations observed in cells downstream of HSC are derived from HSC but are acquired later during the differentiation process.

Finally, we performed gene ontology (GO) enrichment analysis of genes harboring SNVs in both aged murine and human HSCs to identify pathways possibly linked to the positive selection of aged HSCs independent of species. As noted, in aged murine HSCs, we observed significant enrichment for genes involved in epigenetic regulation of gene expression and chromatin organization, cellular organization, metabolism, and transcriptional regulation (p-value < 0.05; **Fig. S2d** and **Table S7**). The two most significant BPs were epigenetic regulation of gene expression and chromatin organization (p < 0.0001; **Fig. S2d** and **Table S7**), biological processes implicated in HSC aging (Buisman and de Haan, 2019; Florian et al., 2018; Grigoryan et al., 2021). Notably, only mutated genes specific to aged stem cells showed enrichment for biological processes, suggesting that the organism selects a specific set of highly related SNVs as it ages. For the human dataset, we observed 3592 and 244 unique age-specific variants and genes, respectively (**Figs. S2** and **S4)**. We also noted enrichment of similar GO terms in the human dataset and, additionally, identified significant enrichment of antigen processing and presentation and of genes related to response to stress and DNA damage (**Fig. S2f**). Changes in epigenetic regulation and chromatin organization linked to SNVs in aged HSCs might thus reflect selection of HSCs during aging, driven by enhanced self-renewal and/or survival over time, in a species-independent manner. Such a view is consistent with the role of epigenetic changes in HSCs during aging.

Taken together, our data refute the notion that an accumulation of mutations is a hallmark of aged HSCs and support a model of clonal selection and expansion of HSCs during aging, with pre-existing aging and MMR mutational signatures. We also report that although the number of clones is similar between young and aged individuals, a subset of clones in aged individuals diverges from the majority. The divergent clones are characterized by a transcriptional and mutational signature reflecting aging, DNA repair deficiency, and downregulation of HSC signature genes. Our data reveal that the mutational and clonal landscape in aged human HSCs is uncoupled from aCH.

## Discussion

We present a simple approach for high-fidelity quantification and evaluation of SNVs in single cells by leveraging cell division biology. In a first approach, both daughter cells arising from the same mother cell were individually sequenced, followed by variant calling and filtering that retained only those present in both daughter cells. Due to the nature of the *ex vivo* model, it primarily identifies homozygous variants in mother HSCs. We conducted both DNA- and RNA-based murine HSC daughter-pair single-cell sequencing and obtained consistent results, indicating a high level of robustness and accurate SNV quantification for this approach. We also performed human HSC single-cell RNA sequencing without daughter-pair information. In this case, we filtered the identified SNVs using a modified version of the daughter-pair analysis model, retaining only those present in at least 2 cells. A Monte Carlo simulation analysis showed that both methods yield high-confidence variants with a near-zero likelihood of such variants occurring by chance. Although the latter approach lacks daughter-cell biology information, it has added value because it inherently includes heterozygous variants. In addition, it is technically less complex, and we were able to sequence more cells per individual, enabling in-depth clonality and deep learning-based aging signature analyses.

We observed a subtle increase in mutation frequency in some aged human HSCs from individual donors compared with young HSCs, but interestingly, the mutational load in murine HSCs remains largely unchanged with age. Thus, although there is an increase in the frequency of DNA mutations in blood cells of the elderly and aged mice (Beerman et al., 2014; Moehrle and Geiger, 2016; Rossi et al., 2007; Xie et al., 2014), our data indicate that mutation frequency is not (in mice) or only slightly (in humans) elevated in aged HSCs. Consequently, the increase in mutation frequency in blood cells upon aging might reflect the acquisition of additional mutations during differentiation, a hypothesis that requires further experimental confirmation. The data are also consistent with findings showing that aged mice, upon irradiation, do not show elevated levels of mutations in hematopoietic cells newly derived from aged HSCs (Moehrle et al., 2015).

The SNV-based clonal evolution analysis revealed a higher level of clonality and genealogical relationships among HSCs from aged mice. In contrast, we did not observe significant differences in clone numbers between young and aged HSCs in both human and mouse samples. Interestingly, in both human and murine datasets, the majority of aged stem cells acquire a mutational signature distinct from that of young HSCs, while a small proportion of aged HSCs present a young-like HSC mutational signature. This aging-associated mutational profile is likely a product of non-uniform accumulation of variants. We observed highly overlapping ontological signatures between murine and human datasets, including epigenetics, where packaging, accessibility, and genome organization of genes bearing aging-associated SNVs are overrepresented. Similar to (Florian et al., 2018; Takayama et al., 2021; Yu et al., 2017), the data imply a strong role for epigenetic configuration, rather than necessarily its transcriptional state, in determining HSC potential. This translates, upon aging, into a more heterogeneous pool of HSC clones with epigenetically defined behaviors. Within human HSCs, we robustly detected SNVs in genes relevant for aCH and thus likely clonal expansion. We detected mutations in these genes in almost all individuals in the study, including young donors. The occurrence of SNVs in these genes is, surprisingly, not specific to aged HSCs. The recurrence of these mutations at the single-cell level was low, with a maximum of ∼10-15% of cells containing one or the other type of mutation, and on average not more than 5% of cells contained one specific mutation. This is, at least in aged individuals, in contrast to what has been reported about the frequency of mutations in genes like *DNMT3A* in BM or blood cells (Jaiswal and Ebert, 2019). Second, the specific aCH mutations we detected in HSCs were different from those found in BM cells from the same donor, which is in contrast to current models of aCH and leukemia, in which the pool of HSCs will contain some of the mutations identified in the downstream pool (Majeti, 2014). This indicates that the genes that are targeted might remain the same in HSCs, though the nature of mutations that are present and which will likely accumulate among HSCs are distinct from the ones that are detected later in BM or blood. This conclusion implies that the primary level of selection of aging-associated human clonal hematopoiesis is not the HSC.

Finally, because mutational signatures reflect the underlying mechanisms by which mutations accumulate (Nik-Zainal et al., 2012), the proportion of aged HSCs sharing similar mutational signatures yet distinct from those of young HSCs reveals an aging signature highly correlated with both age and DNA MMR deficiency. This observation strongly supports the possibility that aged HSCs undergo a time-dependent change in nature, not in mutation load, that increases the likelihood of further evolution into distinct clones, which may then be shaped by selection pressures, as postulated by evolutionary theory (Rozhok et al., 2014). HSCs with mutations in genes involved in epigenetic regulation of gene expression and chromatin organization might possess a survival or proliferative advantage with age. This finding further implies that selection of HSCs with age may not primarily be a consequence of acquiring mutations that make individual HSCs fitter than their young counterparts. Rather, it is an adaptive process within a dynamic, selective evolutionary landscape that also changes with age, shaping the pool of HSCs with specific somatic mutations and clonal competition (Martincorena et al., 2018). Otherwise, young HSCs, which usually show greater stem cell potential than aged HSCs, would simply continue to dominate the HSC pool not only in young individuals but also in aged animals.

## Author Contributions

MAM developed concepts for variant analysis, including custom scripts and hypothesis-validation models; analyzed single-cell data; and wrote the manuscript. MCF and HG designed and performed mouse experiments, generated single-cell sequencing data, and contributed to the manuscript. JH provided support in generating the targeted sequencing dataset. AA, KN, AV, and HG designed and prepared single-cell libraries and generated sequencing data, including human single-cell HSCs. RE, MH, and AL contributed human samples. KB and J-PM performed human single-cell sequencing.

## Acknowledgement

MAM and HG acknowledge the SFB 1506.

## Materials and Methods

### Mice

Young C57BL/6 mice (10-12 weeks old) were obtained from Janvier. Aged C57BL/6 mice (20-26 months old) were obtained from the internal divisional stock (derived from mice obtained from both The Jackson Laboratory and Janvier) as well as from NIA/Charles River. Congenic young and aged C57BL/6.SJL-Ptprc^a^/Boy (BoyJ) mice were obtained from Charles River Laboratories or the internal divisional stock (derived from mice obtained from Charles River Laboratories). All mice were housed in the animal barrier facility under pathogen-free conditions at the University of Ulm or CCHMC. All mouse experiments were performed in compliance with the German Law for Welfare of Laboratory Animals and were approved by the Institutional Review Board of the University of Ulm or by the IACUC of CCHMC.

### Single HSC sorting and culturing

Bone marrow mononuclear cells were flushed from the long bones (tibiae and femurs) of young and aged mice and isolated by low-density centrifugation (Histopaque 1083, Sigma). Low-density bone marrow cells were stained with a cocktail of biotinylated lineage antibodies. The biotinylated antibodies used for lineage staining were all rat anti-mouse antibodies: anti-CD11b (clone M1/70), anti-B220 (clone RA3-6B2), anti-CD5 (clone 53-7.3), anti-Gr-1 (clone RB6-8C5), anti-Ter119, and anti-CD8a (clone 53-6.7) (all from eBioscience). After lineage depletion by magnetic separation (Dynalbeads, Invitrogen), cells were stained with anti-Sca-1 (clone D7) (eBioscience), anti-c-kit (clone 2B8) (eBioscience), anti-CD34 (clone RAM34) (eBioscience), anti-CD127 (clone A7R34) (eBioscience), anti-Flk-2 (clone A2F10) (eBioscience), and Streptavidin (eBioscience). Single long-term hematopoietic stem cells (HSCs, gated as lineage-c-kit+ sca-1+ CD34^-^Flk2^-^) were sorted using a BD FACS Aria III (BD Bioscience) into Terasaki plates. HSCs were microscopically tracked during the culture period. HSCs were cultured in IMDM+10%FBS+P/S+100ng/ml TPO, G-CSF, SCF (PeproTech) at 37°C, 5%CO_2_, 3%O_2_. Each well of the Terasaki plate was microscopically tracked, and within 1 hour of HSC division, daughter cells were separated by mechanical pipetting, deposited into separate Terasaki wells, and quickly rechecked by microscopy. Eventually, the daughter pair underwent scRNA-seq or scATAC-seq preparation.

### Single-cell RNA-seq (SC-RNA-seq)

The SMART-Seq v4 Ultra Low Input RNA kit from Clontech (Catalog # 634892) was used. The kit generates Illumina-compatible RNA-Seq libraries. Sorted single cells were cultured with and without treatment in the presence of cytokines until the first cell division (40-44 hrs). The daughter cells were manually separated, washed with PBS, and collected for RNA sequencing. cDNA synthesis and amplification were performed as recommended by Clontech. The amplified cDNA from single cells was used to prepare libraries with Illumina’s Nextera XT DNA Library Preparation kit (Catalog # FC-131-1096) according to Illumina’s instructions. The libraries were quantified using an Agilent bioanalyzer, pooled, and subjected to next-generation sequencing on Hi-Seq 2500 under paired-end 75-bp conditions.

For the human data set, cells were sorted into 384-well plates containing 1.2 µl of lysis buffer with 0.2% Triton, dNTPs, RNase inhibitor, and the polyT primer, then stored at - 80°C. After thawing, the lysed cells were incubated for 3 min at 72°C, and the reverse transcription mix was added (0.2 U Maxima, 7.5% PEG-8000, RT buffer, TSO, and RNase inhibitor). RT was run for 1,5 h at 42°C, and cDNA was amplified with KAPA HotStart PCR mix and the IS primer. cDNA was purified using Ampure beads, and libraries were prepared in 384-well plates using the Mosquito system with a downscaled version of Illumina’s Nextera XT DNA Library Preparation kit.

### Single-Cell Assay for transposase-accessible chromatin using sequencing (SC-ATAC-Seq)

The protocol for generating ATAC-seq libraries was based on a previously published approach (Buenrostro et al., 2015). Briefly, single HSCs from young and aged C57Bl6 mice were sorted into Terasaki plates with IMDM supplemented with 10% FBS, P/S, 100 ng/ml of SCF, G-CSF, and TPO at 3% O_2_, 5% CO_2_. Microscopically tracked daughter cells obtained after one round of cell division were separated, and each cell was subjected to fragmentation of open chromatin regions using Tn5 transposase (Illumina Inc., San Diego, California, USA) followed by library preparation. The New England Biolabs NEBNext Ultra II Q5 Master Mix (NEB Inc., Ipswich, Massachusetts, USA) was used for the pre-amplification step with the following primers:

Primer 1: 5’-GTCTCGTGGGCTCGGAGATGTGTATAAGAGACAG-3’ and

Primer 2: 5’-TCGTCGGCAGCGTCAGATGTGTATAAGAGACAG-3’.

The generated libraries were quantified using an Agilent bioanalyzer (Santa Clara, California, USA) and a qPCR kit (NEB Inc., Ipswich, Massachusetts, USA). The libraries were then pooled and sequenced on an Illumina HiSeq 3000 (Illumina Inc., San Diego, California, USA) in 75-bp paired-end mode.

**Targeted sequencing of human bone marrow (BM) for** detection of age-related clonal hematopoiesis (aRCH) was performed as described previously (Snetsinger et al., 2019).

### Gene expression quantification and Gene Ontology analysis

Raw Fastq files were adapter-trimmed using Trimm Galore (Babraham Institute). Filtered reads with a Phred score of 20 or higher were aligned to the mouse reference genome version 10 (mm10) using tophat (Trapnell et al., 2009). Transcript abundance estimation, FPKM (Fragments Per Kilobase of transcript per Million mapped reads) calculations, and normalization were performed using Cufflinks (Roberts et al., 2011; Trapnell et al., 2013; Trapnell et al., 2010). Gene Ontology (GO) analysis and biological process enrichment analysis were performed using PANTHER (Mi et al., 2017).

For the human dataset, paired-end raw reads were aligned to the reference genome (Homo_sapiens.GRCh38.99) using HISAT2, and a gene expression count matrix was generated using featureCounts with Homo_sapiens.GRCh38.99.gtf. All downstream analyses were performed using R and Bioconductor (Huber et al., 2015).

### RNA-seq-based variant calling and filtering

Raw reads were trimmed with Trim Galore (Babraham Institute) and aligned to the University of Santa Cruz mouse genome version 10 (UCSC mm10) using the STAR aligner (Dobin et al., 2013). Duplicate marking was performed with Picard tools. Splitting of N Cigar reads, reassigning mapping qualities, and variant calling were performed using the Genome Analysis TK best practices (GATK) (DePristo et al., 2011; Van der Auwera et al., 2013). For subsequent analysis, we excluded low-quality variants and indels.

For the human dataset, we employed UCSC hg19 and followed best-practice variant-calling steps (GATK) (DePristo et al., 2011; Van der Auwera et al., 2013).

### ATAC-seq-based variant calling and filtering

After converting reads to unmapped BAM files, adapter sequences were marked using Picard tools (http://broadinstitute.github.io/picard/). Reads were aligned to the UCSC mm10 genome using Burrows-Wheeler Aligner (bwa) (Trapnell et al., 2009). Duplicate reads were marked using Picard, followed by base recalibration and variant calling with GATK. We retained only quality-filtered single-nucleotide variants for subsequent analyses.

### Targeted sequencing variant calling and filtering

We also applied the GATK best practices for alignment and variant calling (DePristo et al., 2011; Van der Auwera et al., 2013).

### Filtering of variants present in daughter cells (joint filtering)

All post-variant-calling analyses were performed in R and Bioconductor (Huber et al., 2015). Briefly, we first created a single-nucleotide variant (SNV) frame using the VariantAnnotation package (Obenchain et al., 2014). We then used our novel in-house script to apply a joint variant-filtering rule, retaining only variants present in both daughter cells of a given pair. To ensure uniform treatment across all pairs, we further excluded variants that were present in a daughter pair but in only a single member of other pairs.

For the human dataset, because there are no daughter pairs, we considered SNVs present in at least 2 cells per individual for downstream analysis (see the text for further details).

### Monte Carlo Simulation of variant overlaps

After generating empirical overlap information between daughter pairs using bedtools (Quinlan and Hall, 2010), a Monte Carlo Simulation was performed to quantify the chance of randomly encountering such matching events for a given pair. We first generated chromosome-controlled, position-resampled SNVs for each cell using bedtools, then counted overlapping positions. We iterated this process 1000 times for each daughter pair. For each iteration, the number of positional matches was further adjusted by the probability of having the same base substitution at that given position (P = ¼ x ¼ = 0.0625). P-values of the significance of the number of observed overlaps in a given pair were obtained by using the formula 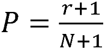, where *r* is the number of iterations in the random overlap analysis that had higher overlap values than the empirical overlap count of that given pair, while *N* is the total number of iterations (1000).

### Deep learning and aging predictors

An R implementation of Keras/TensorFlow (Allaire, 2020) was used to perform binary (young/aged) cross-entropy multilayer deep learning to identify aging predictors from the human single-cell dataset. A 4x16 layer with a glorot uniform initializer and a hyperbolic tangent activation function (tanh) was used to train the model for 500 iterations. Validation on the unseen dataset, along with additional steps including identification of top correlated genes and PCA, was performed using R and Bioconductor (Team, 2015).

### Further Statistical Analyses and Plots

Analyses included distributions of variants across chromosomes, comparisons between age groups, Venn diagrams of overlaps, and associated statistical tests (Wilcoxon test, Chi-squared test, t-test), all performed in the R environment (Team, 2015). A Circos plot was generated using the Perl-based circos package (Krzywinski et al., 2009). Variant presence/absence-based Hamming distance between cells and consensus clustering analysis were performed using the R packages e1071 (Meyer, 2015) and ConsensusClusterPlus (Wilkerson and Hayes, 2010), respectively. Modified heatmaps were plotted using Heatplus (Ploner, 2014). A Neighbor-Joining (NJ)-based phylogenetic tree was generated using Trex (Boc et al., 2012). Mouse-level clonality and variant event ordering analyses were performed using oncoNEM (Ross and Markowetz, 2016). Human clonality analysis was performed using DENDRO (Zhou et al., 2020). Signature analysis was performed using SomaticSignatures (Gehring et al., 2015), while tspair (Leek, 2009) and kernlab (Karatzoglou, 2004) were used to identify top-scoring signature pairs and matching support vector machines (svm), respectively.

