## Supplementary Figures for "A divergent mutational and clonal landscape in aged HSCs is not linked to aging-associated clonal hematopoiesis"

**Fig. S1. (a)** Long term Hematopoietic stem cells (LT-HSC) FACS sorting strategy from young and aged mice **(left panel)** and human **(right panel). (b)** Detailed workflow of the analysis approaches implemented on single-cell RNA and ATAC-seq datasets from mouse and human samples **(c)** ScATAC-seq based variant count in each cell pair before and after applying our filtering rule. Green and blue points represent SNV counts in each daughter cell (A=green, mean ($\bar{x}$) = 1444; B=blue, mean ($\bar{x}$) = 1099; represented by colour matched lines). Purple points indicate the number of position matched variants in daughter pairs (Joint=purple, mean ($\bar{x}$) = 587). The final mean count of position and base substitution matched SNVs is 555. Red points depict the predicted number of positional matches in a Monte Carlo Simulation (MCS) analysis based on position reshuffled daughter pairs (mean ($\bar{x}$) = 0.0047; 1000 iterations). The orange points and line represent the MCS results after adjusting for the probability of base substitution match (mean ($\bar{x}$) = 0.00029). **(d)** Boxplot of single cell distribution of all variants in young and aged HSCs based on scATAC-seq data. Similar to what has been shown in figure 1d (scRNA-seq data), young and aged HSCs show no difference in variant counts across all categories. **(e)** Chromosome size-based plot showing observed and expected variant count across all cells based on the scATAC-seq dataset. Expected frequency was generated by assigning variant count proportional to the size of a given chromosome. The red line depicts the linear relationship between the observed and expected values while the blue line shows the theoretical expected line. Descending order of chromosome sizes is shown on the right side of the plot. **(f)** Chromosome gene density-based plot showing observed and expected variant count across all cells based on the scATAC-seq dataset. Expected frequency was generated by assigning a proportion of total variant count matching the fraction of gene density (number of genes per mega base or genes/Mb) of a given chromosome. The red line depicts the linear relationship between the observed and expected values while the blue line shows the theoretical expected line. Descending order of gene density is shown on the right side of the plot.

**Fig. S2. (a)** Mutated genes overlap between young and aged HSCs in the murine RNA-seq dataset. Coding SNV overlap analysis **(top)** showed that there are 88 variants in common between the age groups. When we then summarized SNVs by genes **(bottom)**, we noted that there are 38 and 31 genes unique to young (green) and aged (blue) HSC, respectively. Fifty-one genes were in common between the age groups. **(b)** Profile of aged HSC specific coding SNVs in the murine dataset. The number of variants per gene is shown by the bar plot in the upper panel. Each rectangle in the lower mutation plot depicts a specific variant in a given cell where the x-axis summarizes the number of variants by the specific cell where they were identified. First y-axis (left) shows a summary of the number of cells where a specific variant was observed and the secondary y-axis shows the gene name associated with every SNV. **(c) (right panel)** Line plot of mean young and aged samples expression levels of the genes with mutations specific to the aged mice. These genes do not show a significant difference in expression between the two groups. **(right panel)** Heatmap showing expression profile of coding SNVs across individual cells. Red and green indicate either higher or lower expression than the mean expression of a gene across all cells, respectively. Group membership of each cell is indicated below the heatmap and bars represent a single cell. There was no significant grouping of young and aged

HSCs in either of the two clusters (Fisher's exact test p-value ~ 0.592; Table S4). **(d)** Radar plot of significantly enriched GO terms based on aged HSC specific mutated genes in the mouse dataset. The scale is $-10\log_{10}P$ where P stands for adjusted p-value. Grey shaded area represents the significance cutoff. Each vertex depicts p-value corresponding to a specific GO term resolution. **(e)** Mutated genes overlap between young and aged HSCs in the human RNA-seq dataset. **(left panel)** We identified 3779 SNVs between the two age groups while 3592 and 2093 were specific to the aged and young LT-HSCs, respectively. **(right panel)** the overlap at the gene level. **(f)** Radar plot of significantly enriched GO terms based on aged HSC specific mutated genes in the human dataset. The scale is $-10\log_{10}P$ where P stands for adjusted p-value and FDR stands for false discovery rate. Grey shaded area represents the significance cutoff. Each blue and red vertex depicts p-value and FDR corresponding to a specific GO term resolution, respectively.

**Fig. S3. (top-left)** The Keras/TensorFlow deep learning analysis approach used to identify an aging signature in the human single-cell RNA-seq dataset. The entire dataset was split into training (65% of cells) and validation set (35%). Model details are provided in the methods section. **(bottom-left)** A biplot showing prediction accuracy using the validation set unseen by the model (~93% accuracy and a chi-squared test of significance of accuracy against a random assignment (50-50) was highly significant with a p-value of 1.88e-66). **(top-right)** Principal component analysis (PCA) using the top 5% predictors based on their correlation to the model. It can be noted that there is a clear segregation between young (green) and aged (blue) LT-HSCs. **(bottom-right)** Publicly available HSC signature and genes upregulated in aging show negative and positive correlation to the genes that are correlated with aged samples in the deep learning model, respectively.

**Fig. S4.** Gene **(top panel)** and SNV **(middle panel)** level overlap between samples in each age group, and, summarized by age groups based on the human single-cell RNA-seq dataset. Bottom panel shows the overlap between aged-specific mutated genes in the human and mouse dataset and only two genes (*LUC7L* and *SRRM1*) were identified. Both of these genes are splicing regulators.

**Fig. S5.** Representative clonality plots of a young and an aged sample. After identifying the optimal k cluster based on a screeplot (number of clusters after which a further increment in cluster number does not yield reduction in intragroup sum of squared estimate of errors, SSE), shown here are number of identified clones along with number of cells per clone, mutation count per clone, distance between clones, a phylogram of individual cells colored by their clone membership, and a similarity heatmap based on hamming distance between individual cells using clonality-relevant mutations.

**Fig. S6. (a)** The residuals sum of squares (RSS; upper panel) and proportion of variance explained (lower panel) based on a principal component analysis of 40 somatic signatures generated using the coding SNVs in the mouse dataset. The first three signatures explain ~ 50% of the variance in the data while half of the generated signatures (20) explain 90% variation. **(b)** Principal component analysis (PCA) based Mutational signature profile of the mouse dataset. The six types of substitutions are shown on top while bases flanking each type of substitution are given at the bottom. Y-axis (left axis) indicates the percentage of contribution of each motif (a combination of a given substitution and the associated flanking bases). Only the signatures of interest (S5 and S13) are shown. **(c)** A Venn diagram showing overlap between significant PCA mutational signatures based on Odds-Ratio (OR) and the result of top scoring pair (tsp) analysis in the mouse dataset. The OR analysis (purple circles) indicate the number of signatures that have significant OR in the class assignment of young and aged HSCs (29 signatures at $p < 0.05$ and 11 at $p < 0.01$). The blue circle indicates the two signatures with the highest classification score in the tsp analysis. It can be noted that there is a full overlap between the tsp and OR analyses. A non-negative matrix factorization (nmf) based ven diagram showing overlap between significant nmf mutational signatures based on an Odds-Ratio (OR) and top scoring pair (tsp) analyses. The OR analysis (purple circles) indicate the number of signatures that have significant OR in the class assignment of young and aged HSCs (34 signatures at $p < 0.05$ and 27 at $p < 0.01$). The blue circle indicates the three signatures with the highest classification score in the tsp analysis. It can be noted that there is full overlap between the tsp and OR analyses. **(d)** A contour plot showing subgroups of young and aged HSCs based on support vector machine (svm) analysis of S5 and S13 in the mouse dataset. Blue contours represent aged HSCs while green depicts young HSCs. The white area indicates the boundary between the two classes. We noted five and two clusters for the young and aged HSCs, respectively (white circles with dashed lines), indicating higher heterogeneity in aged HSCs. (e) Correlation between published somatic signatures with known biological association (blue labels) and the three nmf based signatures of the current dataset (S12, S26, and S35; shown in red labels). Only the statistically significant correlations ($p < 0.05$) are shown, represented by the blue ellipses whose color intensities and shape show the strength of correlation). The overlaid purple box highlights the significant correlation between S26, S35, and the published aging and MMR signatures.

**Fig. S7.** Coding SNVs based evolutionary relationship among cells (cellular/clonal evolution) in all mice (6 young and 6 Aged) in the study based on the scRNA-seq dataset. Cells are placed in the hierarchy based on maximum likelihood-based estimation based on a nested structure of the mutational pattern. Cells not linked with arrows indicate that there is no sufficient deducible relationship based on the provided SNV data. It can be noted that half of the aged HSC mice show a simple proliferation like the hierarchical link between cells while the young HSC show little or no intercellular evolutionary relationship. The heatmap show profile of false positive rate (FPR, x-axis) and false negative rate (FNR, y-axis) in a given analysis, the inverted triangle within each plot indicates the estimated FPR/FNR for that analysis.

**Fig. S8. (top-left panel)** Proportion of variance explained based on a principal component analysis (PCA) of 10 somatic signatures of representative young and aged samples. **(right panel)** Top scoring pair (tsp) and support vector machine (svm) analyses of all possible pairs of combinations between 10 signatures in the aged and young sample. After grouping the cells into two groups primarily based expression analysis and on the first component of a between group analysis (bga; see upper and lower left panels in Figure 4b and Fig. S9), shown here are pairs ranked according to their cross-validation error rate in a svm analysis (the best combination being the ones with the lowest error rate). **(bottom-left panel)** a contour biplot of top signatures of the representative mutational signatures. It can be noted that the aged cells were clearly separated into two groups while the young cells did not show clear segregation.

**Fig S9. (a)** Clonality within a representative young sample and clonal grouping-based gene expression analysis. We detected three clones in this sample. A pairwise differential expression followed by Hamming's distance between the pairs of comparisons reveals the most distant set is clone 1 to clone 3. The rest of the pairs had high similarity (blue part of heatmap), except another pair of comparison including clone 3. A 3D principal component analysis (PCA) using original group coloring (top part; 3 colors) and a recolored one based on the pairwise differential expression analysis (lower part; 2 colors where clone 3 is colored blue while the other clones are colored red). We also performed GSEA (right panel) where we show hematopoietic stem cell and DNA damage signature show significant upregulation in the clones 1 and 2 as compared to clone 3. We also performed GSEA of the top 5% predictors from the deep learning analysis and could show that they show upregulation in the clones 1 and 2 indicating that clone 3 is not only genetically but also transcriptional more distinct from the other clones. **(b)** A non-negative matrix factorization (nmf) mutational signatures matrix based between groups analysis (bga) shows that clone 3 is more distinct from the other clones **(left panel)**, in line with what we showed with the transcriptional analysis. We then tested 10 signatures in a pairwise fashion (using top scoring pairs and support vector machines, svm) to identify the pairs with the highest discrimination score. Here is shown the svm plot **(middle panel)** of signatures 6 and 10, where clone 3 in blue shows clear segregation from the rest of the clones (red). **(right panel)** a correlation between signatures 4 and 8 and publicly available signatures with known biological association. We observe here, in the case of a young sample, that there is no correlation between our signatures and the published ones.

**Fig. S10. (top panel)** Clonal hematopoiesis (aCH) gene expression profile in young and aged HSCs in the human single-cell RNA-seq dataset. We did not observe significant difference between the age groups and all genes showed median expression levels that are higher than the global median expression profiles. This is critical, as a proof of principle, that the mutations detected (or not detected) are not confounded by insufficient coverage or differences between age groups. (**bottom left panel**) Mutations detected in the aCH genes analyzed shown here both at single-cell level and sample level. (**bottom right panel**) Heatmap and barplot showing recurrence of individual mutations across age groups and their frequency across sample, respectively.

**Fig. S11.** Coverage specific to the 11 aCH genes based on targeted sequencing panel. Fraction of capture target bases by depth plot (left) and heatmap of target and immediate flanking regions (right) show coverage across genes of interest, with coverage ranging from 250 – 3500, with a median coverage of ~1500.

**Fig. S12.** Gene level summary of mutation recurrence, number of unique mutations per gene, and allelic frequency of mutations are shown in the left and middle panel. Right panel shows recurrence of individual mutations across the 6 samples sequenced (blue bar above the plot = Aged; green bar above the plot = Young).

**Fig. S13. (a)** SNV count across cell types (HSCs have the lowest median SNV count). **(b)** Hierarchical clustering of cell types based on SNV overlap. **(c)** Correspondence analysis (two dimensions) showing similarity between samples based on types of SNVs. **(d)** Circos plot (highlighting HSCs in red) showing relative number of shared SNVs between cell types.

**Supplementary Tables:**

**Table S1.** The frequency of SNVs detected in the single-cell RNA-seq and single-cell ATAC-seq dataset. Age group, cell pair ID, individual variant count, overlaps, adjusted overlaps, mean predicted overlaps (simulated), and adjusted mean predicted overlaps (r = counts the number of instances in the simulation where variant frequencies higher than the empirical counts are observed). *(Worksheet "Supplementary_table_1" in the attached excel file "Supplementary_tables.xlsx")*

**Table S2.** Wilcoxon test on the significance of the difference in SNV frequency between young and aged HSCs. We generally observed no significant difference between the age groups before and after applying our joint variant calling algorithm. *(Worksheet "Supplementary_table_2" in the attached excel file "Supplementary_tables.xlsx")*

**Table S3.** Expression levels of mutated genes. We did not see a significant difference between young and aged HSCs (see Benjamini-Hochberg adjusted p-value), indicating that expression levels of do not contribute towards differential detection (specificity) of SNVs to a given age group compared to the other. *(Worksheet "Supplementary_table_3" in the attached excel file "Supplementary_tables.xlsx")*

**Table S4.** Group membership (unsupervised hierarchical clustering) of samples based on the expression levels of the mutated genes. In line with our results shown in Table S3, we saw a random assignment of samples into two groups, with no significant overrepresentation of either young or aged HSCs in a given group (Fisher's exact test; p-value not significant). *(Worksheet "Supplementary_table_4" in the attached excel file "Supplementary_tables.xlsx")*

**Table S5.** List of mutated genes specific to aged HSCs. Similar to the large gene list presented under Table S3, we also confirmed here that the specificity of these genes does not depend on their higher expression levels in the aged group (none of the adj. p-values are significant). *(Worksheet "Supplementary_table_5" in the attached excel file "Supplementary_tables.xlsx")*

**Table S6.** Based on the mutated genes specific to aged HSCs, we also saw a random assignment of samples into two groups, with no significant overrepresentation of an age group in a given cluster (Fisher's exact test; p-value not significant). *(Worksheet "Supplementary_table_6" in the attached excel file "Supplementary_tables.xlsx")*

**Table S7.** Panther Gene Ontology (GO) terms with significant enrichment based on mutated genes specific to aged HSCs. Of note, neither the young HSC specific nor genes overlapping between young and aged HSCs showed significant enrichment. *(Worksheet "Supplementary_table_7" in the attached excel file "Supplementary_tables.xlsx")*

**Table S8.** Binary matrix (presence/absence) of SNVs (young HSCs) used as input in the clonality analysis. Mouse ids are shown on top, followed by cell ids. SNVs are shown in the first column. *(Worksheet "Supplementary_table_8" in the attached excel file "Supplementary_tables.xlsx")*

**Table S9.** The binary matrix of SNVs used as input in the clonality analysis of aged HSC. *(Worksheet "Supplementary_table_9" in the attached excel file "Supplementary_tables.xlsx")*

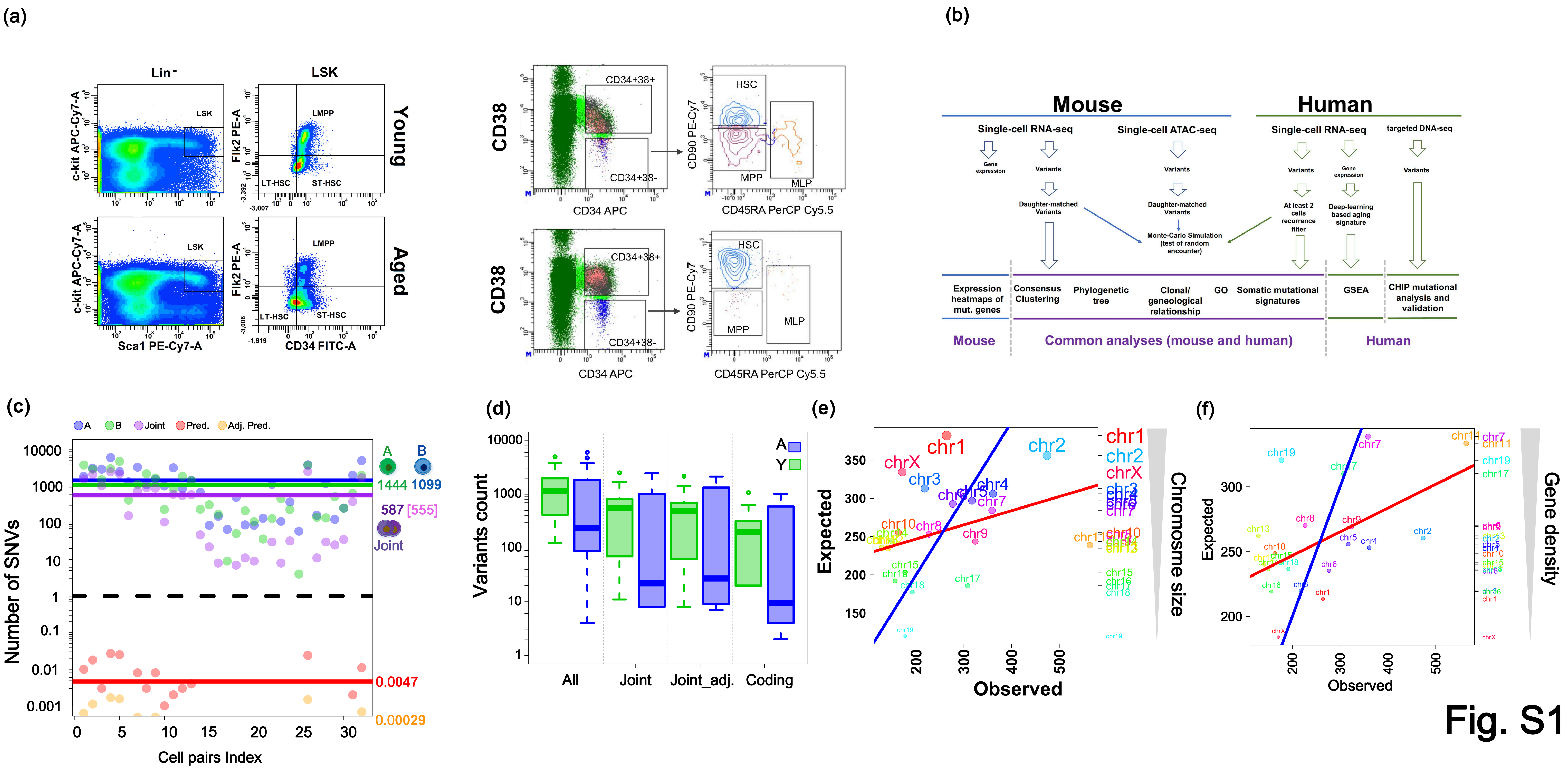

(a)
Lin-
LSK
Young
Aged
c-kit APC-Cy7-A
Sca1 PE-Cy7-A
Flk2 PE-A
CD34 FITC-A
LMPP
LT-HSC
ST-HSC
CD38
CD34 APC
CD34+38+
CD34+38-
CD90 PE-Cy7
CD45RA PerCP Cy5.5
HSC
MPP
MLP
(b)
Mouse
Human
Single-cell RNA-seq
Single-cell ATAC-seq
targeted DNA-seq
Gene expression
Variants
Daughter-matched Variants
Monte-Carlo Simulation (test of random encounter)
At least 2 cells recurrence filter
Deep-learning based aging signature
Expression heatmaps of mut. genes
Consensus Clustering
Phylogenetic tree
Clonal/ geneological relationship
GO
Somatic mutational signatures
GSEA
CHIP mutational analysis and validation
Mouse
Common analyses (mouse and human)
Human
(c)
A
B
Joint
Pred.
Adj. Pred.
Number of SNVs
Cell pairs Index
1444
1099
587 [555]
0.0047
0.00029
(d)
Variants count
All
Joint
Joint_adj.
Coding
(e)
Expected
Observed
Chromosome size
(f)
Expected
Observed
Gene density


Fig. S1

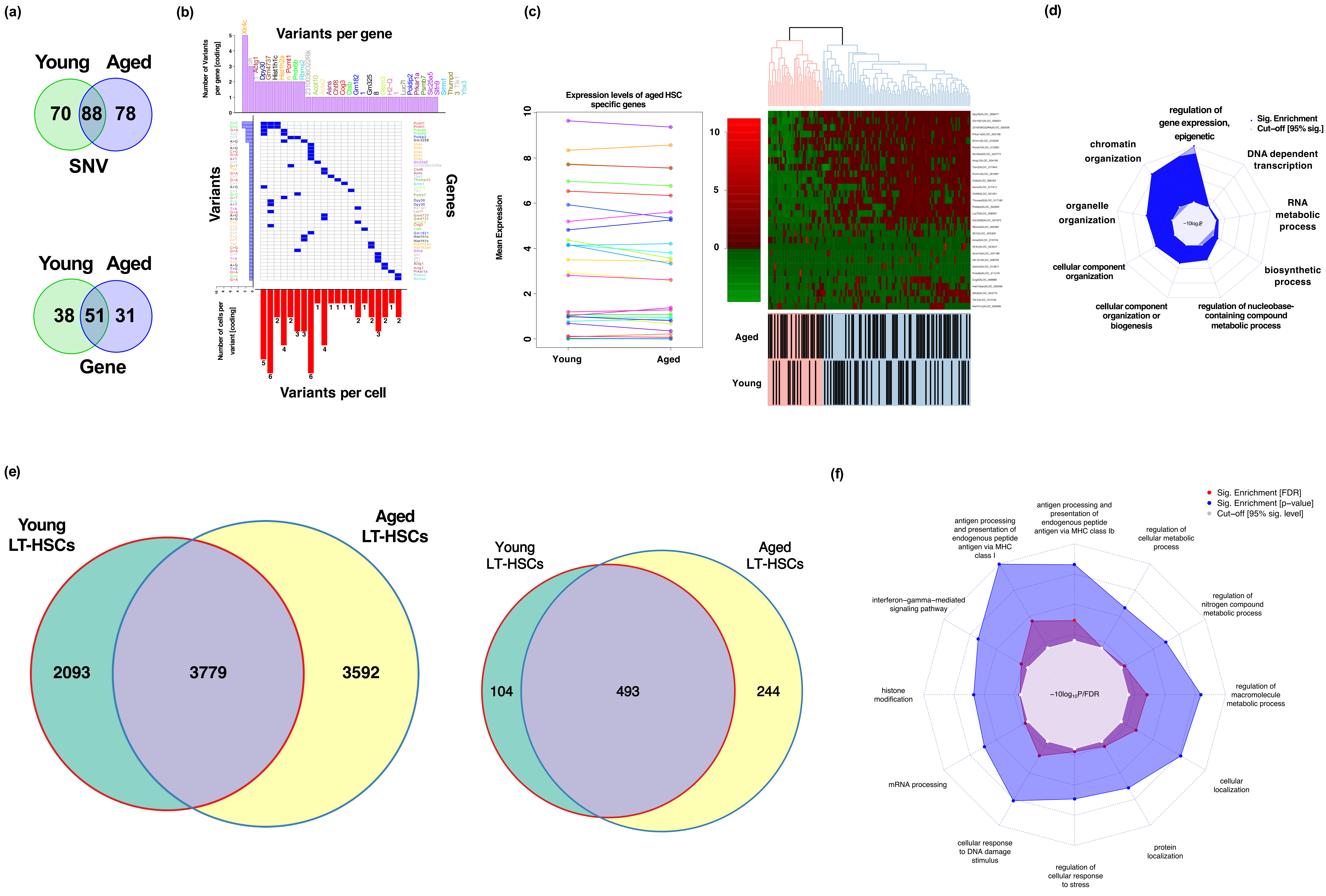
(a)
Young Aged
70 88 78
SNV
Young Aged
38 51 31
Gene
(b)
Variants per gene
Number of Variants per gene [coding]
Variants
Genes
Number of cells per variant [coding]
Variants per cell
(c)
Expression levels of aged HSC specific genes
Mean Expression
Young
Aged
10
5
0
Aged
Young
(d)
regulation of gene expression, epigenetic
Sig. Enrichment
Cut−off [95% sig.]
chromatin organization
DNA dependent transcription
organelle organization
RNA metabolic process
cellular component organization
biosynthetic process
cellular component organization or biogenesis
regulation of nucleobase-containing compound metabolic process
(e)
Young LT-HSCs
Aged LT-HSCs
2093
3779
3592
Young LT-HSCs
Aged LT-HSCs
104
493
244
(f)
Sig. Enrichment [FDR]
Sig. Enrichment [p−value]
Cut−off [95% sig. level]
antigen processing and presentation of endogenous peptide antigen via MHC class Ib
antigen processing and presentation of endogenous peptide antigen via MHC class I
regulation of cellular metabolic process
interferon−gamma−mediated signaling pathway
regulation of nitrogen compound metabolic process
histone modification
regulation of macromolecule metabolic process
mRNA processing
cellular localization
cellular response to DNA damage stimulus
protein localization
regulation of cellular response to stress
−10log10P/FDR

Fig. S2

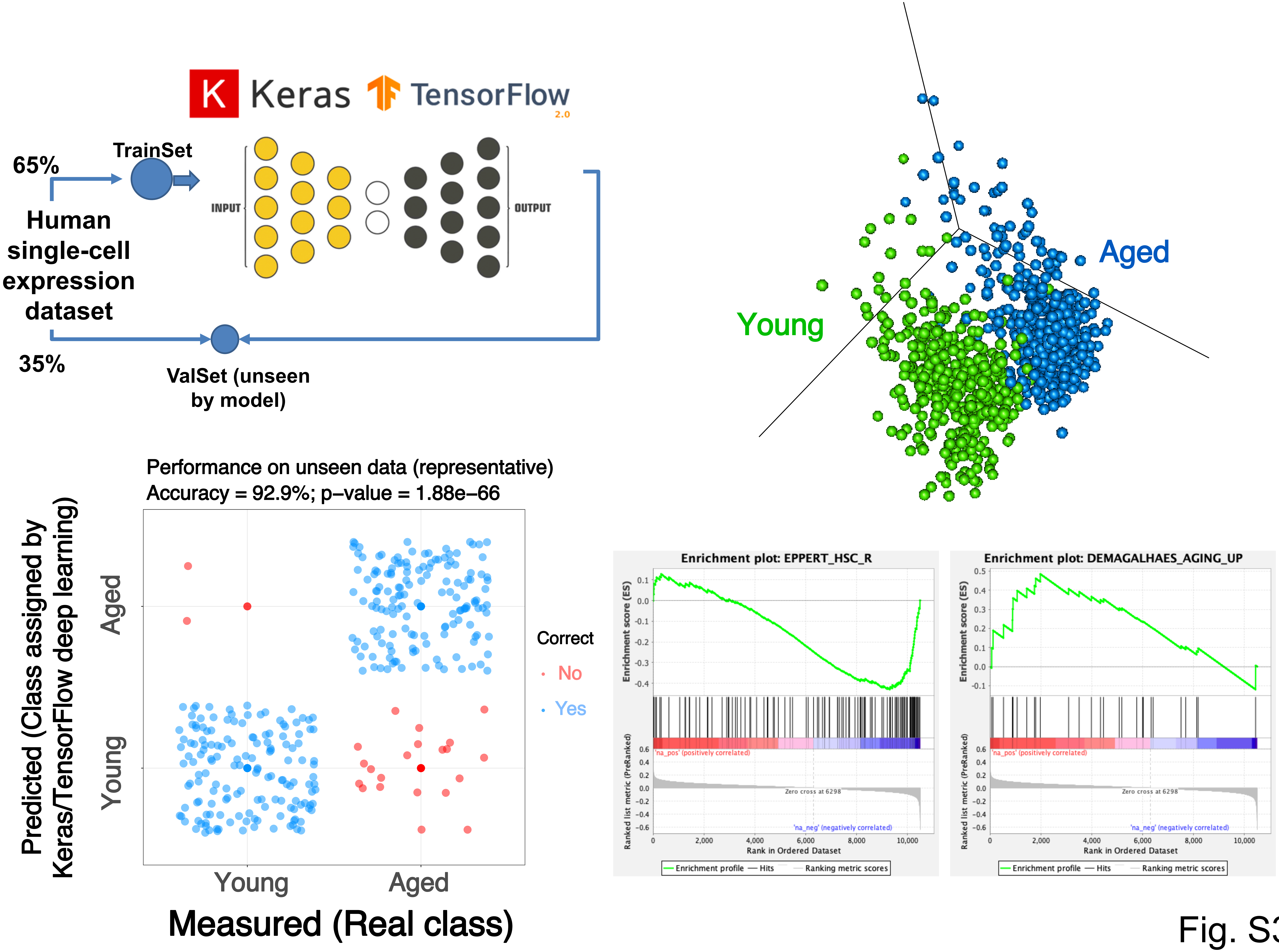
Keras
TensorFlow
2.0
65%
TrainSet
Human single-cell expression dataset
INPUT
OUTPUT
35%
ValSet (unseen by model)
Aged
Young
Performance on unseen data (representative)
Accuracy = 92.9%; p–value = 1.88e–66
Predicted (Class assigned by Keras/TensorFlow deep learning)
Aged
Young
Correct
No
Yes
Young
Aged
Measured (Real class)
Enrichment plot: EPPERT_HSC_R
Enrichment score (ES)
Ranked list metric (PreRanked)
'na_pos' (positively correlated)
Zero cross at 6298
'na_neg' (negatively correlated)
Rank in Ordered Dataset
Enrichment profile
Hits
Ranking metric scores
Enrichment plot: DEMAGALHAES_AGING_UP
Enrichment score (ES)
Ranked list metric (PreRanked)
'na_pos' (positively correlated)
Zero cross at 6298
'na_neg' (negatively correlated)
Rank in Ordered Dataset
Enrichment profile
Hits
Ranking metric scores

Fig. S3

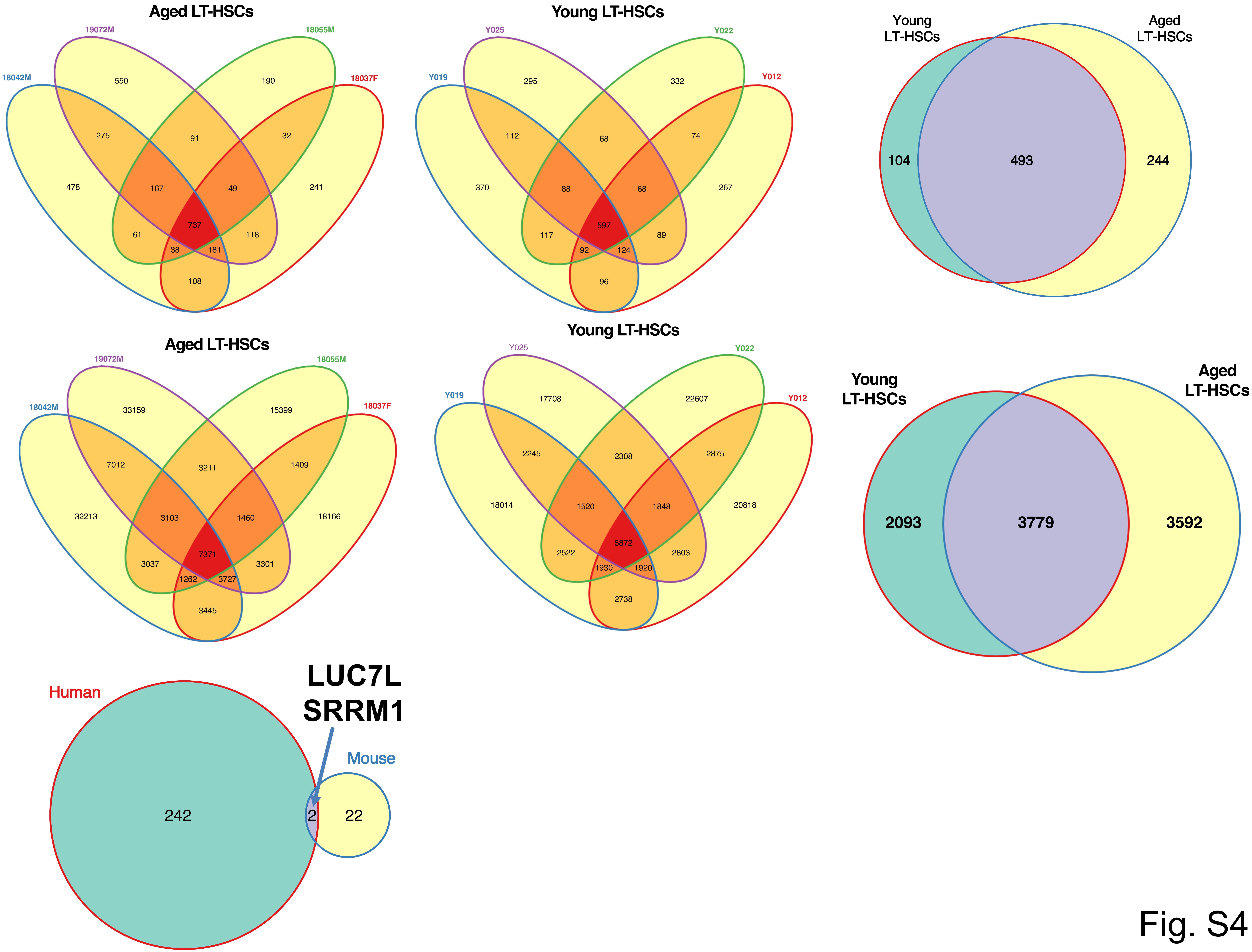
Aged LT-HSCs
19072M
18055M
18042M
18037F
550
190
275
91
32
478
167
49
241
737
61
118
38
181
108
Young LT-HSCs
Y025
Y022
Y019
Y012
295
332
112
68
74
370
88
68
267
597
117
89
92
124
96
Young
LT-HSCs
Aged
LT-HSCs
104
493
244
Aged LT-HSCs
19072M
18055M
18042M
18037F
33159
15399
7012
3211
1409
32213
3103
1460
18166
7371
3037
3301
1262
3727
3445
Young LT-HSCs
Y025
Y022
Y019
Y012
17708
22607
2245
2308
2875
18014
1520
1848
20818
5872
2522
2803
1930
1920
2738
Young
LT-HSCs
Aged
LT-HSCs
2093
3779
3592
Human
LUC7L
SRRM1
Mouse
242
2
22

Fig. S4

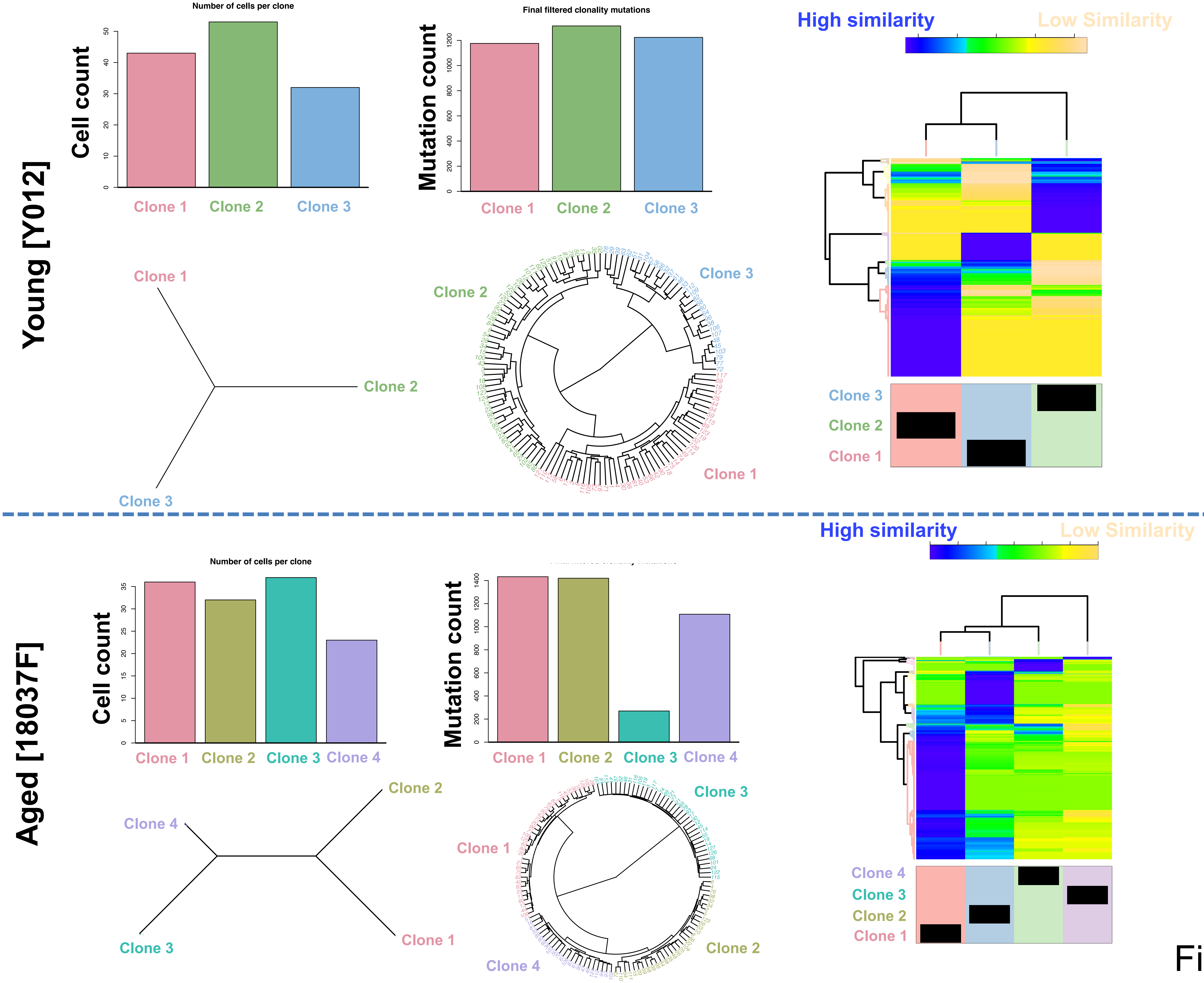
Young [Y012]
Number of cells per clone
Cell count
Clone 1
Clone 2
Clone 3
Final filtered clonality mutations
Mutation count
Clone 1
Clone 2
Clone 3
High similarity
Low Similarity
Clone 1
Clone 2
Clone 3
Clone 2
Clone 3
Clone 1
Clone 3
Clone 2
Clone 1
Aged [18037F]
High similarity
Low Similarity
Number of cells per clone
Cell count
Clone 1
Clone 2
Clone 3
Clone 4
Mutation count
Clone 1
Clone 2
Clone 3
Clone 4
Clone 2
Clone 4
Clone 3
Clone 1
Clone 3
Clone 1
Clone 2
Clone 4
Clone 4
Clone 3
Clone 2
Clone 1

Fig. S5

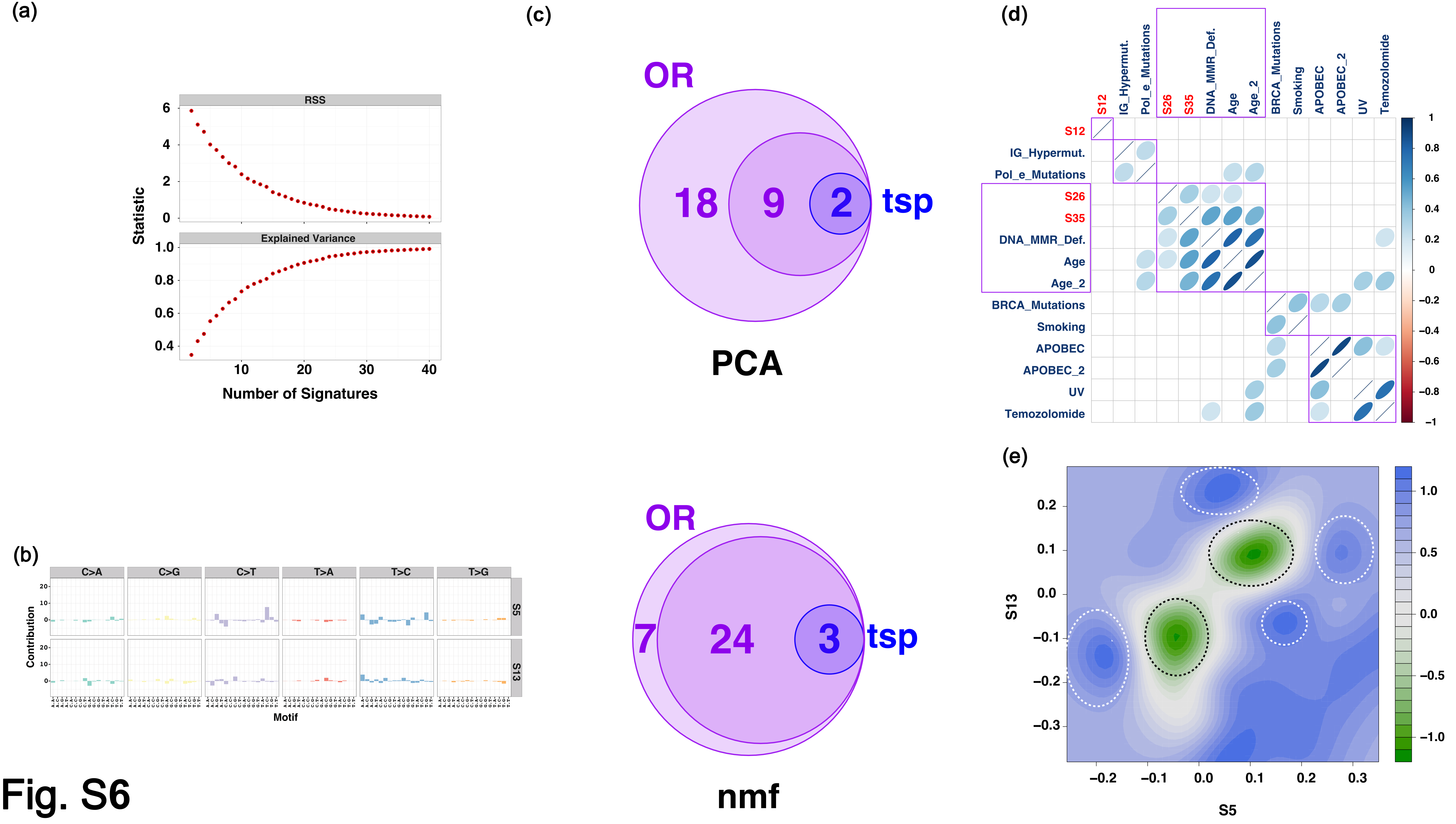
(a)
RSS
Explained Variance
Statistic
Number of Signatures
(b)
C>A
C>G
C>T
T>A
T>C
T>G
S5
S13
Contribution
Motif
(c)
OR
18
9
2
tsp
PCA
OR
7
24
3
tsp
nmf
(d)
S12
IG_Hypermut.
Pol_e_Mutations
S26
S35
DNA_MMR_Def.
Age
Age_2
BRCA_Mutations
Smoking
APOBEC
APOBEC_2
UV
Temozolomide
(e)
S13
S5

Fig. S6

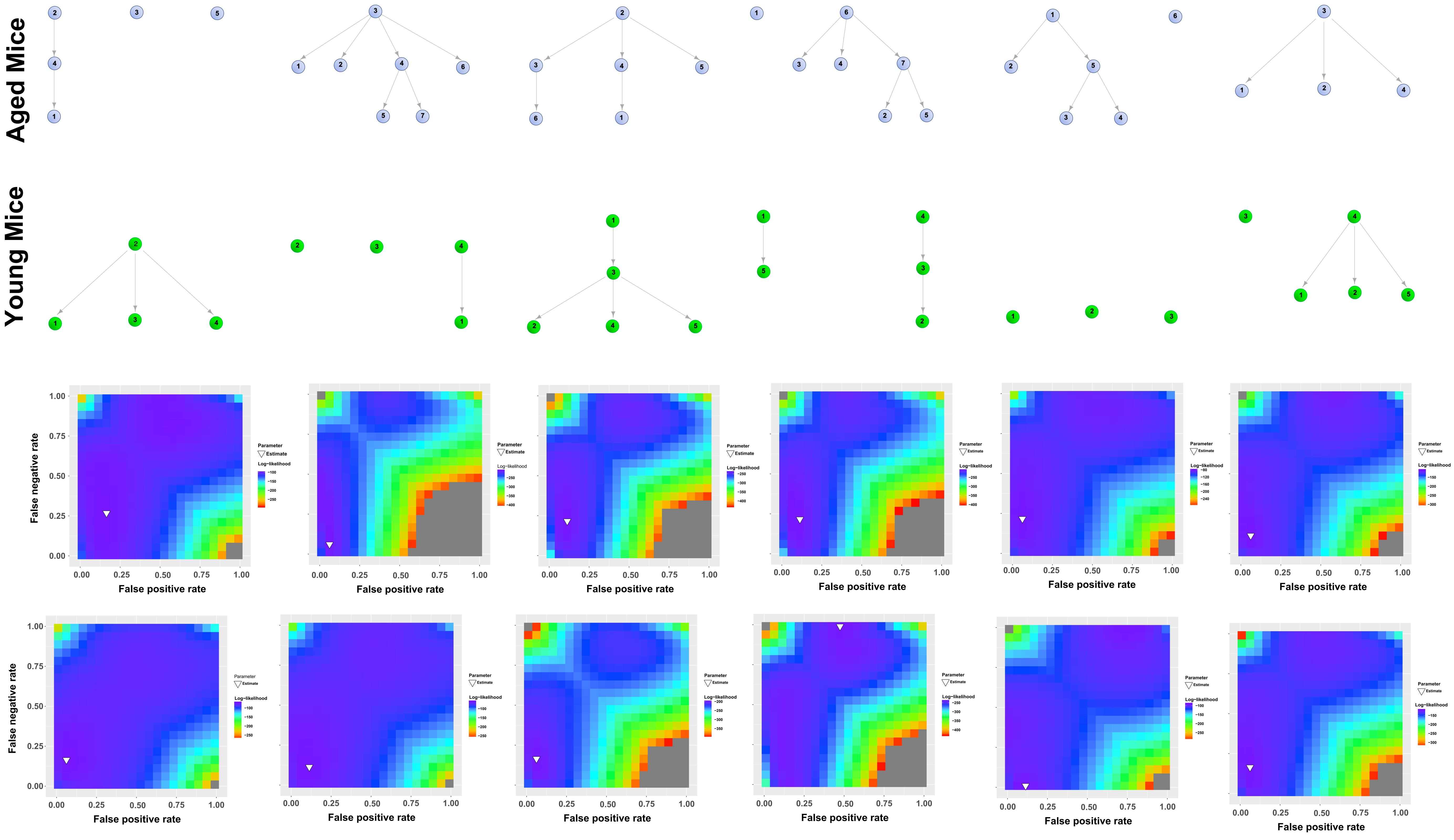
Aged Mice
Young Mice
False negative rate
False positive rate
0.00
0.25
0.50
0.75
1.00
Parameter
Estimate
Log−likelihood

Fig. S7

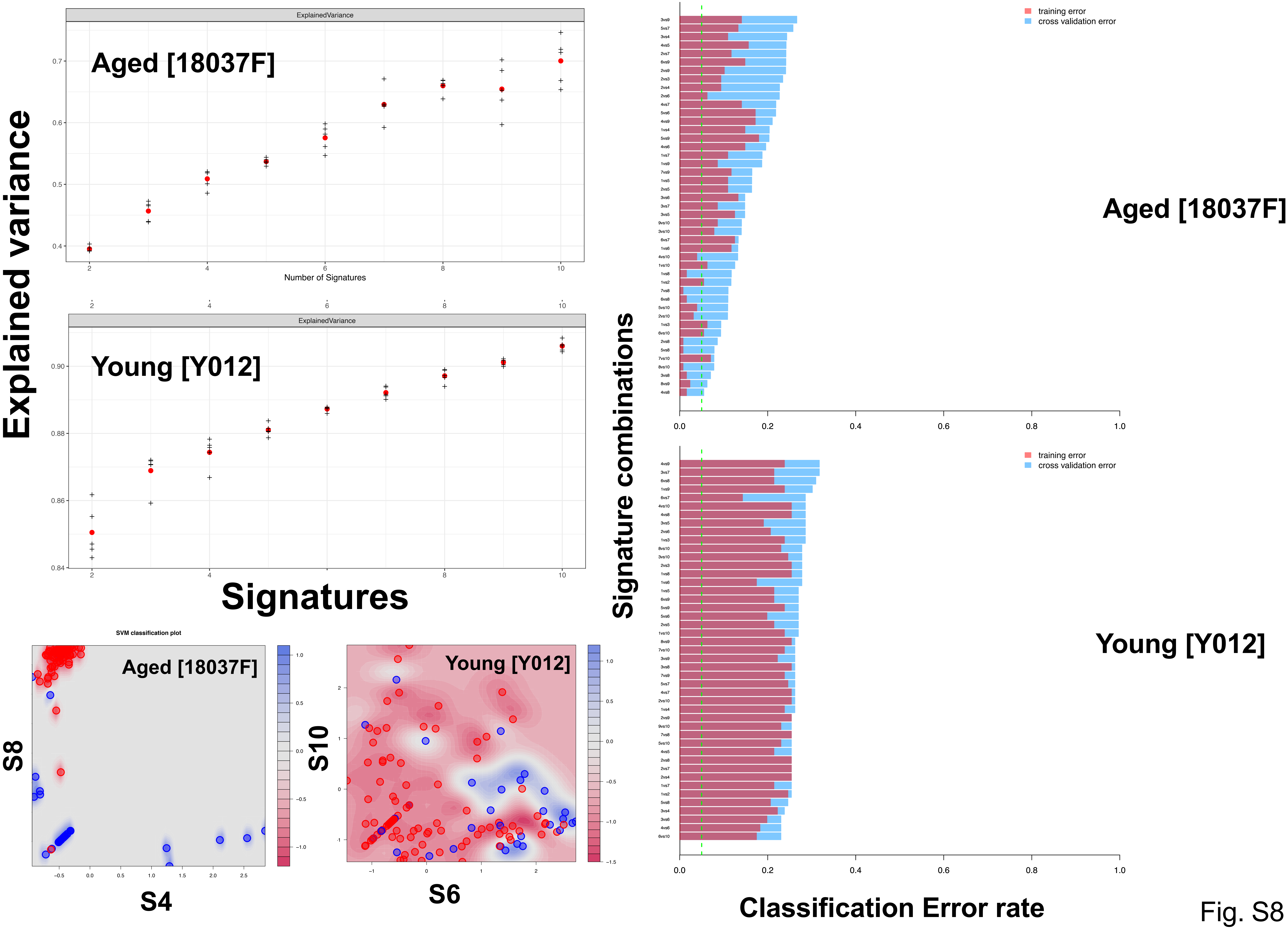

Explained variance
ExplainedVariance
Aged [18037F]
Number of Signatures
Young [Y012]
Signatures
SVM classification plot
S8
S4
S10
S6
Signature combinations
training error
cross validation error
Classification Error rate


Fig. S8

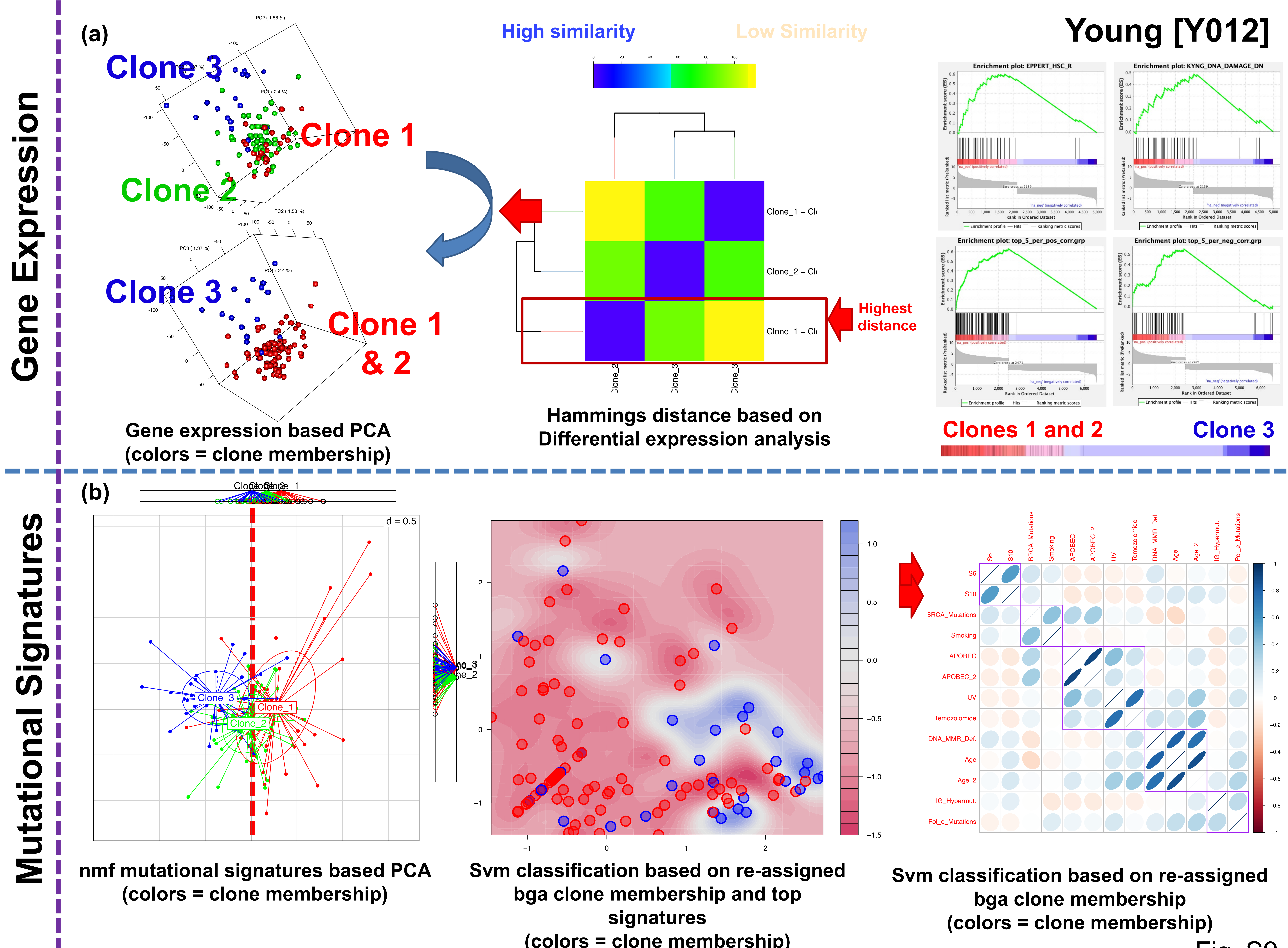
Young [Y012]
Gene Expression
(a)
Clone 3
Clone 1
Clone 2
Clone 3
Clone 1 & 2
Gene expression based PCA (colors = clone membership)
High similarity
Low Similarity
Highest distance
Hammings distance based on Differential expression analysis
Enrichment plot: EPPERT_HSC_R
Enrichment plot: KYNG_DNA_DAMAGE_DN
Enrichment plot: top_5_per_pos_corr.grp
Enrichment plot: top_5_per_neg_corr.grp
Clones 1 and 2
Clone 3
Mutational Signatures
(b)
d = 0.5
Clone_3
Clone_1
Clone_2
nmf mutational signatures based PCA (colors = clone membership)
Svm classification based on re-assigned bga clone membership and top signatures (colors = clone membership)
S6
S10
BRCA_Mutations
Smoking
APOBEC
APOBEC_2
UV
Temozolomide
DNA_MMR_Def.
Age
Age_2
IG_Hypermut.
Pol_e_Mutations
Svm classification based on re-assigned bga clone membership (colors = clone membership)

Fig. S9

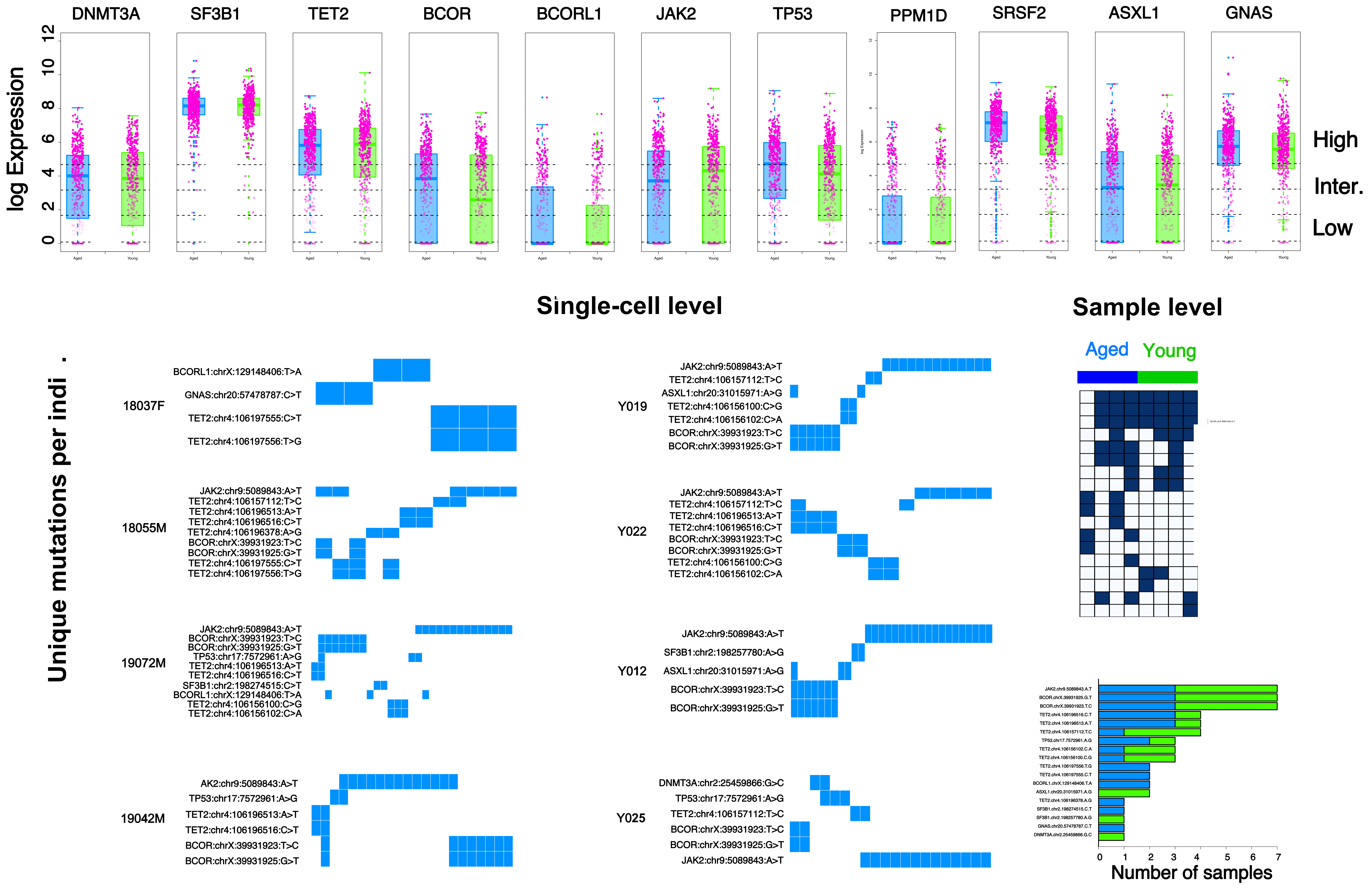

DNMT3A
SF3B1
TET2
BCOR
BCORL1
JAK2
TP53
PPM1D
SRSF2
ASXL1
GNAS
log Expression
0
2
4
6
8
10
12
High
Inter.
Low
Single-cell level
Sample level
Aged
Young
Unique mutations per indi .
18037F
BCORL1:chrX:129148406:T>A
GNAS:chr20:57478787:C>T
TET2:chr4:106197555:C>T
TET2:chr4:106197556:T>G
18055M
JAK2:chr9:5089843:A>T
TET2:chr4:106157112:T>C
TET2:chr4:106196513:A>T
TET2:chr4:106196516:C>T
TET2:chr4:106196378:A>G
BCOR:chrX:39931923:T>C
BCOR:chrX:39931925:G>T
TET2:chr4:106197555:C>T
TET2:chr4:106197556:T>G
19072M
JAK2:chr9:5089843:A>T
BCOR:chrX:39931923:T>C
BCOR:chrX:39931925:G>T
TP53:chr17:7572961:A>G
TET2:chr4:106196513:A>T
TET2:chr4:106196516:C>T
SF3B1:chr2:198274515:C>T
BCORL1:chrX:129148406:T>A
TET2:chr4:106156100:C>G
TET2:chr4:106156102:C>A
19042M
AK2:chr9:5089843:A>T
TP53:chr17:7572961:A>G
TET2:chr4:106196513:A>T
TET2:chr4:106196516:C>T
BCOR:chrX:39931923:T>C
BCOR:chrX:39931925:G>T
Y019
JAK2:chr9:5089843:A>T
TET2:chr4:106157112:T>C
ASXL1:chr20:31015971:A>G
TET2:chr4:106156100:C>G
TET2:chr4:106156102:C>A
BCOR:chrX:39931923:T>C
BCOR:chrX:39931925:G>T
Y022
JAK2:chr9:5089843:A>T
TET2:chr4:106157112:T>C
TET2:chr4:106196513:A>T
TET2:chr4:106196516:C>T
BCOR:chrX:39931923:T>C
BCOR:chrX:39931925:G>T
TET2:chr4:106156100:C>G
TET2:chr4:106156102:C>A
Y012
JAK2:chr9:5089843:A>T
SF3B1:chr2:198257780:A>G
ASXL1:chr20:31015971:A>G
BCOR:chrX:39931923:T>C
BCOR:chrX:39931925:G>T
Y025
DNMT3A:chr2:25459866:G>C
TP53:chr17:7572961:A>G
TET2:chr4:106157112:T>C
BCOR:chrX:39931923:T>C
BCOR:chrX:39931925:G>T
JAK2:chr9:5089843:A>T
0
1
2
3
4
5
6
7
Number of samples


Fig. S10

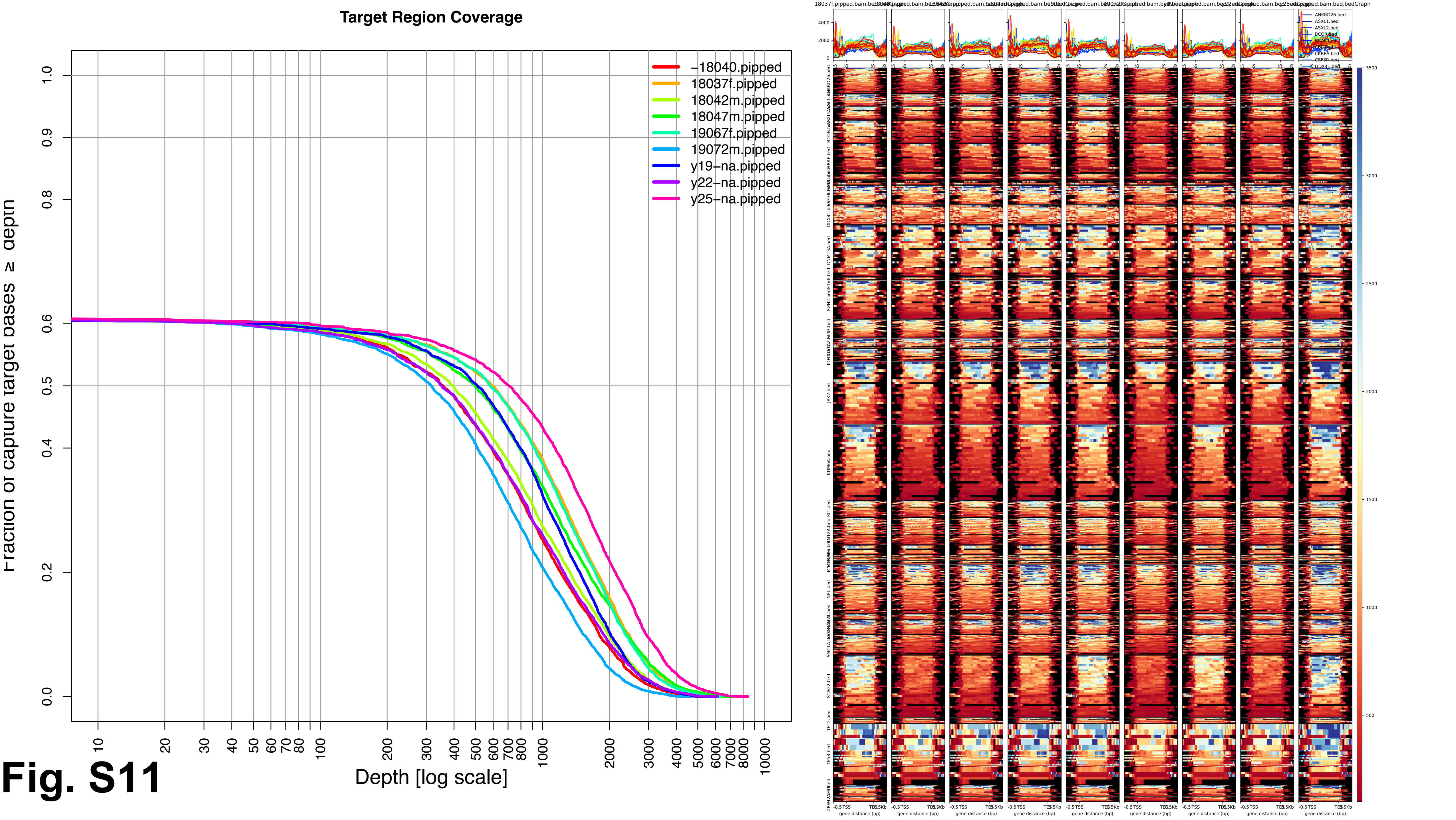
Target Region Coverage
Fraction of capture target bases ≥ depth
Depth [log scale]
-18040.pipped
18037f.pipped
18042m.pipped
18047m.pipped
19067f.pipped
19072m.pipped
y19-na.pipped
y22-na.pipped
y25-na.pipped
0.0
0.2
0.4
0.5
0.6
0.8
0.9
1.0
10
20
30
40
50
60
70
80
100
200
300
400
500
600
700
800
1000
2000
3000
4000
5000
6000
7000
8000
10000
gene distance (bp)
JAK2.bed
KDM6A.bed
STAG2.bed
TET2.bed
TP53.bed
NF1.bed
DNMT3A.bed
3500
3000
2500
2000
1500
1000
500

Fig. S11

### Fig. S12

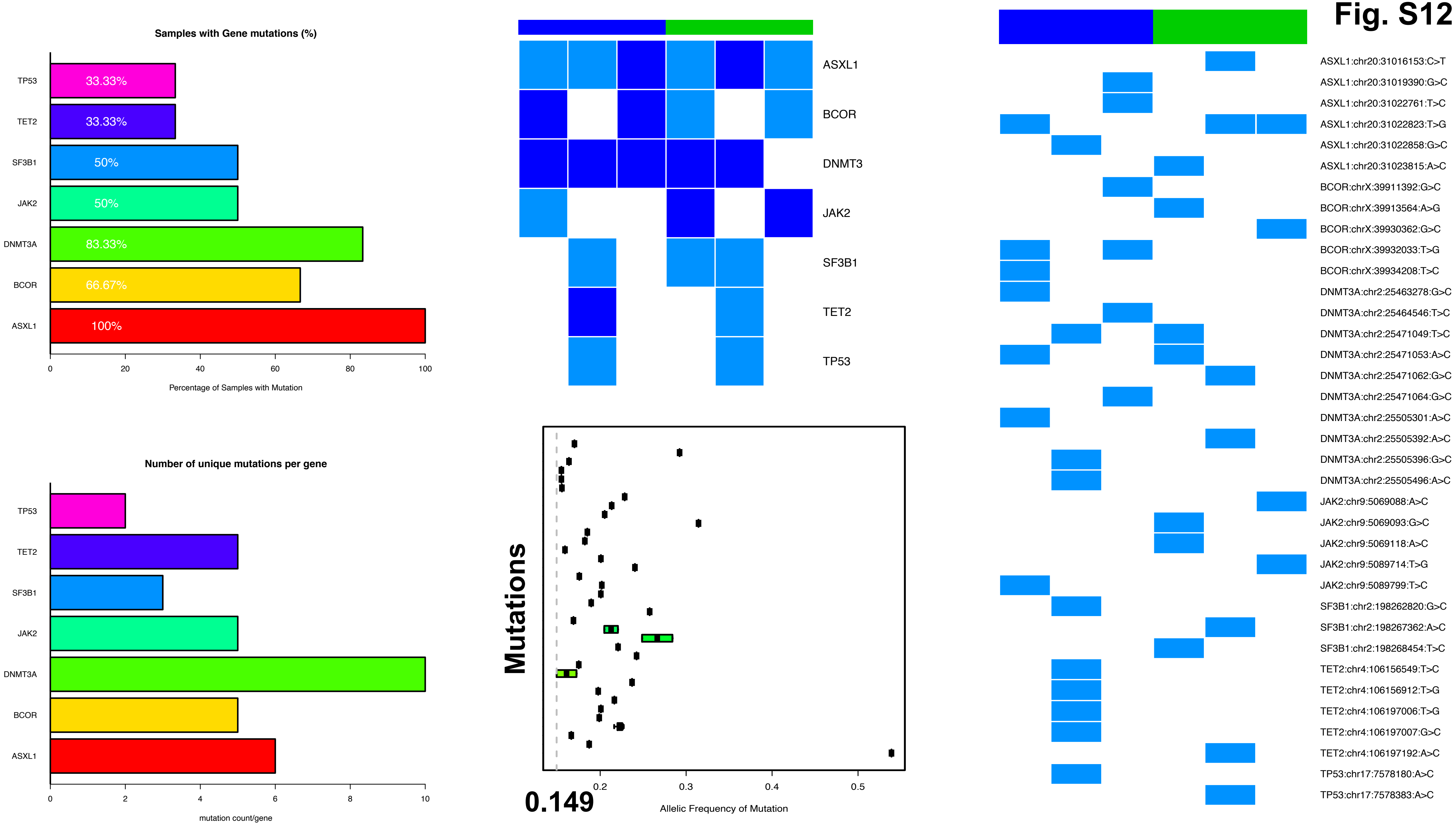

Samples with Gene mutations (%)
TP53
TET2
SF3B1
JAK2
DNMT3A
BCOR
ASXL1
33.33%
33.33%
50%
50%
83.33%
66.67%
100%
0
20
40
60
80
100
Percentage of Samples with Mutation
Number of unique mutations per gene
0
2
4
6
8
10
mutation count/gene
ASXL1
BCOR
DNMT3
JAK2
SF3B1
TET2
TP53
Mutations
0.149
0.2
0.3
0.4
0.5
Allelic Frequency of Mutation
ASXL1:chr20:31016153:C>T
ASXL1:chr20:31019390:G>C
ASXL1:chr20:31022761:T>C
ASXL1:chr20:31022823:T>G
ASXL1:chr20:31022858:G>C
ASXL1:chr20:31023815:A>C
BCOR:chrX:39911392:G>C
BCOR:chrX:39913564:A>G
BCOR:chrX:39930362:G>C
BCOR:chrX:39932033:T>G
BCOR:chrX:39934208:T>C
DNMT3A:chr2:25463278:G>C
DNMT3A:chr2:25464546:T>C
DNMT3A:chr2:25471049:T>C
DNMT3A:chr2:25471053:A>C
DNMT3A:chr2:25471062:G>C
DNMT3A:chr2:25471064:G>C
DNMT3A:chr2:25505301:A>C
DNMT3A:chr2:25505392:A>C
DNMT3A:chr2:25505396:G>C
DNMT3A:chr2:25505496:A>C
JAK2:chr9:5069088:A>C
JAK2:chr9:5069093:G>C
JAK2:chr9:5069118:A>C
JAK2:chr9:5089714:T>G
JAK2:chr9:5089799:T>C
SF3B1:chr2:198262820:G>C
SF3B1:chr2:198267362:A>C
SF3B1:chr2:198268454:T>C
TET2:chr4:106156549:T>C
TET2:chr4:106156912:T>G
TET2:chr4:106197006:T>G
TET2:chr4:106197007:G>C
TET2:chr4:106197192:A>C
TP53:chr17:7578180:A>C
TP53:chr17:7578383:A>C

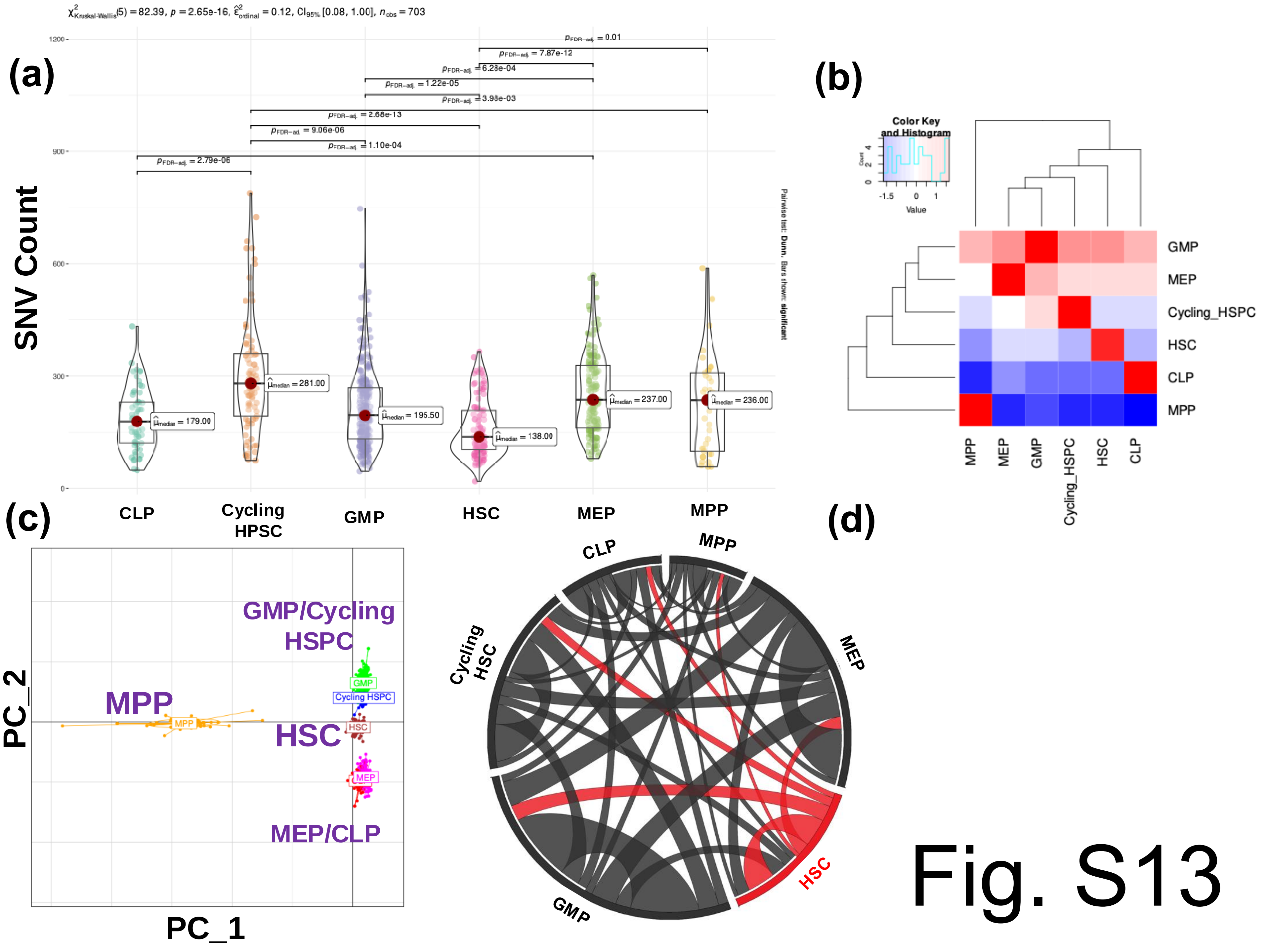
(a)
SNV Count
CLP
Cycling HPSC
GMP
HSC
MEP
MPP
$\hat{\mu}_{median}$ = 179.00
$\hat{\mu}_{median}$ = 281.00
$\hat{\mu}_{median}$ = 195.50
$\hat{\mu}_{median}$ = 138.00
$\hat{\mu}_{median}$ = 237.00
$\hat{\mu}_{median}$ = 236.00
$p_{FDR-adj.}$ = 2.79e-06
$p_{FDR-adj.}$ = 1.10e-04
$p_{FDR-adj.}$ = 9.06e-06
$p_{FDR-adj.}$ = 2.68e-13
$p_{FDR-adj.}$ = 3.98e-03
$p_{FDR-adj.}$ = 1.22e-05
$p_{FDR-adj.}$ = 6.28e-04
$p_{FDR-adj.}$ = 7.87e-12
$p_{FDR-adj.}$ = 0.01
Pairwise test: Dunn, Bars shown: significant
(b)
Color Key and Histogram
Value
GMP
MEP
Cycling_HSPC
HSC
CLP
MPP
(c)
PC_2
PC_1
GMP/Cycling HSPC
MPP
HSC
MEP/CLP
(d)
CLP
MPP
MEP
Cycling HSC
GMP
HSC

Fig. S13
